# The Hypoxia-regulated Ectonucleotidase CD73 is a Host Determinant of HIV Latency

**DOI:** 10.1101/2022.08.03.502655

**Authors:** Hannah S. Sperber, Kyle A. Raymond, Mohamed S. Bouzidi, Tongcui Ma, Silvana Valdebenito, Eliseo Eugenin, Nadia R. Roan, Steven G. Deeks, Sandra Winning, Joachim Fandrey, Roland Schwarzer, Satish K. Pillai

## Abstract

Deciphering the mechanisms underlying viral persistence is critical to achieving a cure for HIV infection. We implemented a systems approach to discover molecular signatures of HIV latently-infected CD4+ T cells, identifying the immunosuppressive, adenosine-producing ectonucleotidase CD73 as a key surface marker of latent cells. Hypoxic conditioning, reflecting the lymphoid tissue microenvironment, increased the frequency of CD73+ CD4+ T cells and promoted HIV latency. Transcriptomic profiles of CD73+ CD4+ T cells favored viral quiescence, immune evasion, and cell survival. CD73+ CD4+ T cells were capable of harboring a functional HIV reservoir and reinitiating productive infection *ex vivo*. CD73 or adenosine receptor blockade facilitated latent HIV reactivation *in vitro*, mechanistically linking adenosine signaling to viral quiescence. Finally, tissue imaging of lymph nodes from HIV-infected individuals on antiretroviral therapy revealed spatial association between CD73 expression and HIV persistence *in vivo*. Our findings warrant development of HIV cure strategies targeting the hypoxia-CD73-adenosine axis.

## INTRODUCTION

In recent decades, remarkable scientific and biomedical advances have turned the tides in the ongoing fight against the global human immunodeficiency virus (HIV) epidemic. However, current therapeutic regimens fail to completely eradicate HIV due to the persistence of latently-infected cells^1^. These viral reservoirs are established very early during infection, remain invisible to the host’s immune system, and persevere for decades despite effective antiretroviral therapy (ART)^2–4^. Although latently-infected cells are extremely rare^5–7^, they are able to reinvigorate spreading infection rapidly, and people living with HIV (PLWH) almost inevitably experience viral rebound within weeks of a treatment interruption^2,8^.

The identification of reliable biomarkers or unique gene expression patterns in HIV latently-infected cells are key goals of current HIV research efforts. Such factors could contribute to the development of a cure by: 1) refining and broadening our understanding of HIV latency mechanisms and the biology of viral persistence, 2) enabling accurate quantification of viral reservoirs to assess viral burden and efficacy of therapeutic interventions, and 3) providing potential therapeutic targets to specifically eliminate viral sanctuaries. However, the heterogenous and dynamic nature of the viral reservoir greatly complicates this endeavor^9^.

Viral reservoirs are found in a variety of anatomical sites and cell types. Infected CD4+ T cells arguably constitute the most important HIV reservoir^10^, and within this highly diverse cell lineage, the expression of several cellular factors is associated with increased levels of integrated proviral DNA^11^. This includes immune checkpoint molecules such as programmed cell death protein 1 (PD-1), cytotoxic T-lymphocyte-associated protein 4 (CTLA-4), lymphocyte-activation gene 3 (LAG-3), and T cell immunoreceptor with Ig and ITIM domains (TIGIT)^9,12,13^. Moreover, expression levels of CD2 in CD4+ T cells were reported to identify HIV latently-infected cells^14^, while CD20^15^ and CD30^16^ expressing CD4+ T cells were found to be specifically enriched for HIV RNA. In 2017, Descours et al.^17^ proposed CD32a as a viral reservoir marker and described an unprecedented 1000-fold enrichment in HIV DNA in CD32a+ cells as compared to CD32a-CD4+ T cells. However, this finding has been repeatedly challenged in subsequent studies and incited a controversial discussion^18–25^.

In the present study, we sought to provide a better understanding of the phenotypic nature of latently infected CD4+ T cells and the mechanisms involved in the establishment and maintenance of HIV reservoirs. To address this aim, we utilized a dual-reporter HIV construct that enables isolation and purification of uninfected, productively-infected, and latently-infected primary CD4+ T cells by flow cytometry^26,27^. We then characterized each of these purified cell populations using systems approaches to obtain comprehensive gene and protein expression profiles of HIV latently-infected cells. Observed molecular signatures of latency were validated using *ex vivo* mechanistic experiments as well as profiling of tissue samples from HIV-infected individuals on suppressive ART. Our data reveal a novel mechanism of HIV latency establishment and maintenance, identifying the hypoxia-CD73-adenosine signaling axis as a key target for therapeutic intervention and diagnostic evaluation in the setting of HIV cure.

## RESULTS

### Sorting of HIV_DFII_-infected primary CD4+T cells enables isolation of latently-infected cells

Profiling of HIV latently-infected cells has been greatly impeded by their extremely low frequencies *in vivo* in HIV-infected individuals, as well as the lack of an established marker enabling purification of latent cells for downstream characterization. To characterize latent HIV reservoir cells, we utilized a modified version of the HIV Duo-Fluo II (HIV_DFII_) single round, recombinant HIV dual-reporter virus (Figure S1A). To achieve sufficient infection frequencies, we first activated blood-derived primary CD4+T cells obtained from six healthy donors *in vitro* via αCD3/αCD28 bead stimulation and infected them with HIV_DFII_ via spinoculation (Figure S1B-D). Four days post infection (p.i.), productively- and latently-infected, as well as uninfected cells were purified by fluorescence activated cell sorting (FACS, Figure 1A). Expectedly^27^, low frequencies of latently-infected cells were established in all donors, whereas uninfected and productively-infected cells could be rapidly collected by FACS in large numbers (Figure 1B and C). After testing the sorted samples for sufficient enrichment of the desired cell populations (Figure S1E), we subjected them, together with a panel of control specimens (untreated samples; unstimulated, infected cells; and stimulated cells without HIV_DFII_ infection) to downstream analysis using systems approaches (Figure 1A). All samples were characterized using mass cytometry by time of flight (CyTOF), measuring 40 surface proteins to provide immunophenotypic profiles at the single cell level. In parallel, we applied NanoString hybridization and fluorescence-based digital counting technology allowing for simultaneous detection of 770 mRNA and 30 protein targets.

**Figure 1:**
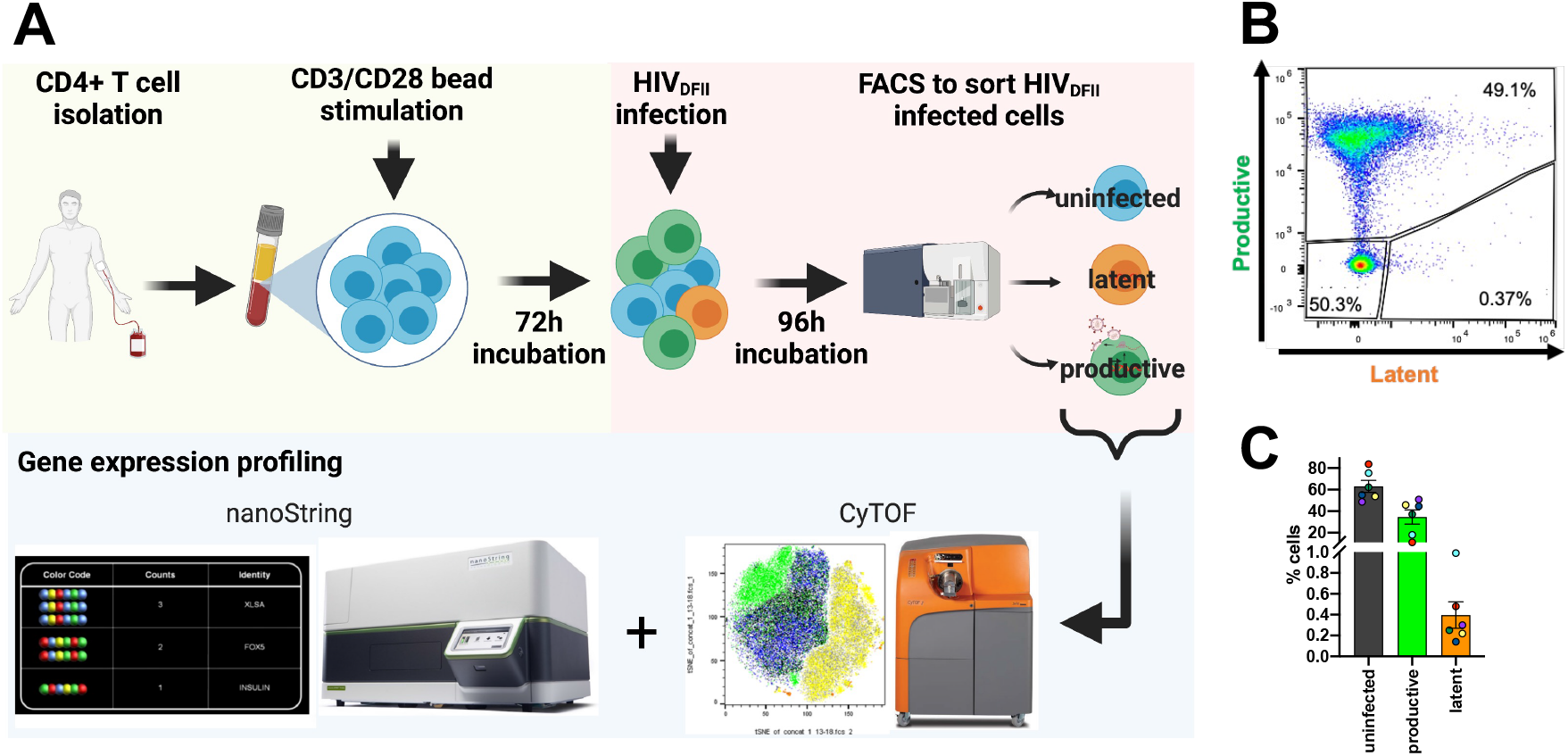
Isolation of latently-infected primary CD4+ T cells using the dual-reporter virus HIV_DFII_. (A) Experimental workflow: Blood-derived primary CD4+ T cells were isolated from six healthy donors and stimulated *in vitro* with αCD3/αCD28 beads for three days. Cells were then infected with HIV_DFII_ and subjected to FACS 4 days post infection. (B) Representative gating for FACS of productively-infected, latently-infected, and uninfected CD4+ T cells. All samples were pre-gated for live, single cells. (C) Bar graph represents the frequencies of infected cells in all six donors. Colored dots indicate individual donors. Error bars show standard error of the mean (SEM).

### Latently-infected cells are phenotypically diverse and include T_regs_ and Tfh cells

First, to establish an in-depth analysis of the phenotypic features of latently-infected cells, we implemented CyTOF, simultaneously quantifying the expression levels of 40 different proteins with single-cell resolution. Our labeling panel comprised T cell lineage and differentiation markers, activation markers, homing receptors, and several proteins that were described previously in the context of HIV latency^9,28,29^ (Table S1). We performed extensive high dimensional analysis and generated t-distributed stochastic neighbor embedding (tSNE) plots to visualize the data and assess specific T cell subsets and population phenotypes. Across all donors, we identified latent cells in various T cell compartments, including central memory (Tcm, CD45RO+CD45RA-CCR7+CD27+), follicular helper (Tfh, CD45RO+CD45RA-PD1+CXCR5+), regulatory (Treg, CD45RO+CD45RA-CD127-CD25+), and to a lesser extent, naïve (CD45RO-CDRA+CCR7+) T cells (Figure 2). Although memory CD4+ T cell subset frequencies exhibited variability between donors and no significant associations with latency were identified, Treg cells were most commonly represented in latent cell populations. Interestingly, CD30, a previously reported marker of transcriptionally-active reservoir cells^16^, was expressed on a subset of latent cells in all analyzed donors (Figure 2).

**Figure 2:**
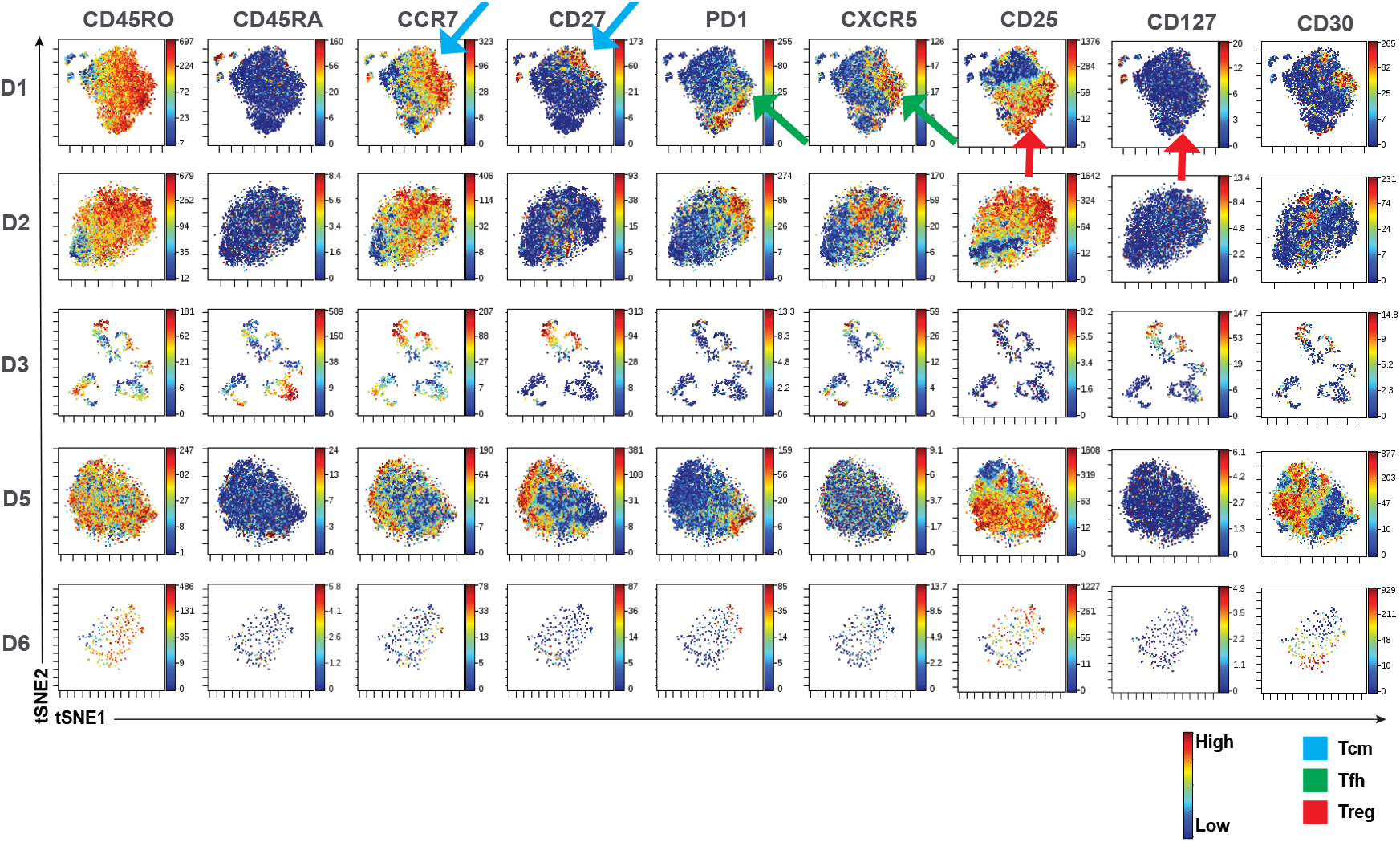
Multiple T cell subsets including T_regs_ are represented among HIV latently-infected cells. (A) Shown are tSNE plots of CyTOF data describing sorted latent cells from each of 5 donors (D1-D6); each donor is analyzed within its own tSNE space. tSNEs are color-coded according to expression levels of the antigen listed at the top (with red corresponding to highest expression, and blue the lowest). Most cells exhibited a memory phenotype, defined by high levels of CD45RO and low levels of CD45RA expression (first two columns). Latent cells exhibited phenotypic features of Tcm cells (CCR7+CD27+), Tfh cells (PD1+CXCR5), and T_regs_ (CD25+CD127-). Latent cells expressing CD30 (last column), a previously-described marker of reservoir cells, were observed in all donors. Areas of the tSNE corresponding to these three cell subsets are highlighted by the blue, green, and red arrows, respectively, for D1.

### NanoString analysis reveals unique features of latently-infected cells that promote viral quiescence and cell survivorship

We next obtained high-dimensional NanoString data to comprehensively characterize gene and protein expression patterns of latent cells. Our initial unsupervised clustering analysis revealed that donor effects dominated expression patterns (Figure 3A, see dendrogram). Importantly, despite substantial donor effects, we observed that expression patterns in uninfected and productively-infected cells within each donor clustered with respect to latent cells, suggesting that latently-infected cells exhibited distinct expression signatures (Figure 3A, see dendrogram).

**Figure 3:**
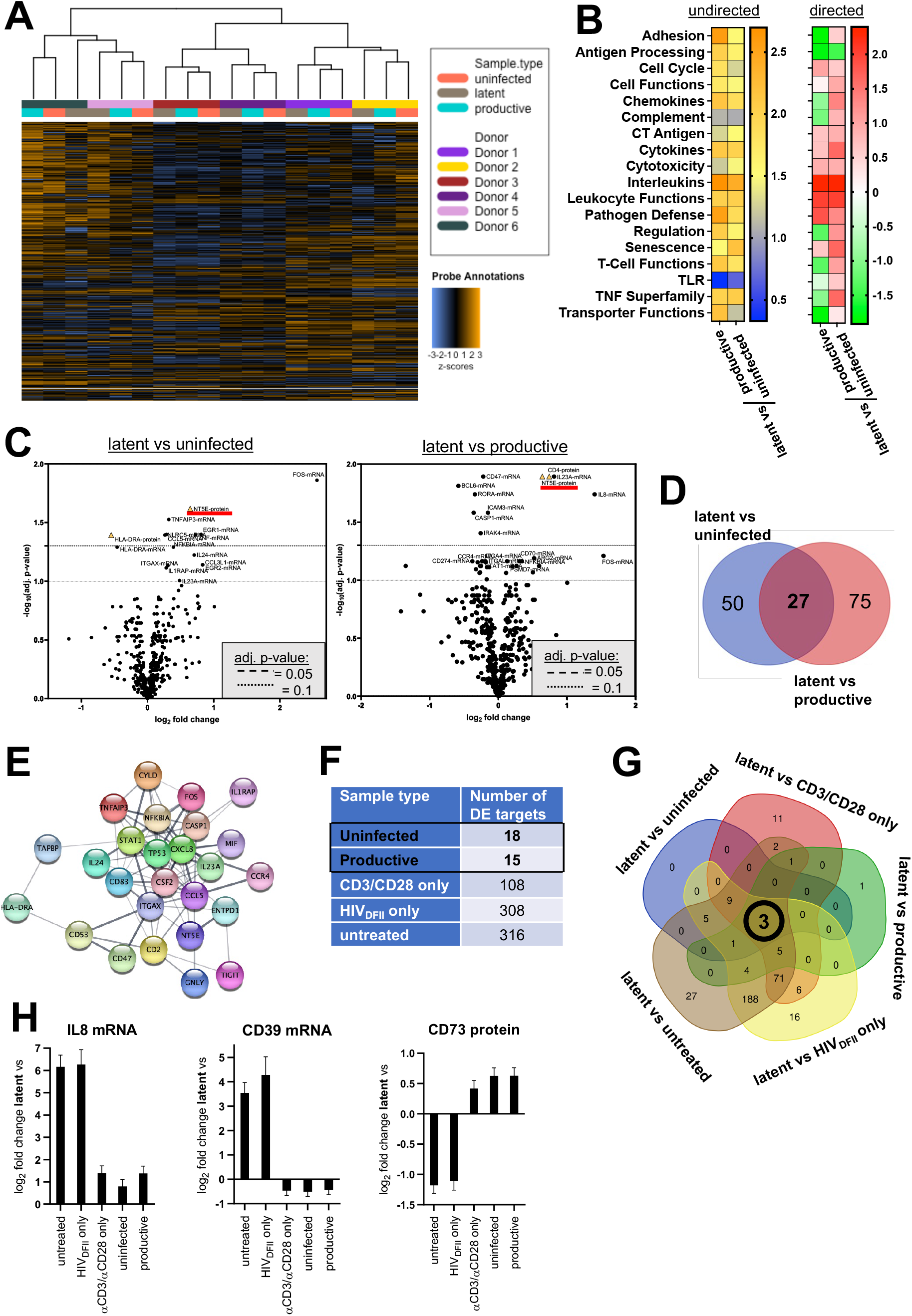
Latently-infected cells share signature features including distinct expression of IL8 mRNA, CD39 mRNA and CD73 protein. (A) Heatmap visualizing normalized gene expression data from unsupervised clustering analysis of virus-exposed, sorted samples. Each row shows normalized counts of a single probe (target) and each column represents an individual sample. Colored horizontal bars along the top identify assigned sample attributes (donor or sample type, see color legend). Hierarchical clustering was used to generate dendrograms. (B) Heatmaps showing undirected and directed global significance scores for signaling pathways based on differential gene expression analysis. The level of significance is indicated from blue to orange, the up- and down-regulation of pathways from red to green. (C) Volcano plots depict differential mRNA (circle) and protein (triangle) target expression between latent vs uninfected and latent vs productively-infected cells. Each data point represents one target. The log_2_ fold change is displayed on the x-axis and the - log_10_ of the adjusted (adj.) p-value (using the Benjamini-Hochberg method) on the y-axis. Horizontal lines indicate adj. p-value thresholds. The top 20 hits in each comparison are labeled. Hits were considered significant at an adj. p-value cutoff < 0.1. NT5E (CD73) is underlined in red. (D) Venn diagram illustrating overlapping hits from the DE analyses ‘latent vs uninfected’ and ‘latent vs productive’ using a p-value cutoff p < 0.05. (E) A protein-protein interaction network of targets uniquely changed in latent cells was generated in Cytoscape 3.8.2. The STRING database for Homo sapiens was applied with a confidence score of 0.4. Thickness of lines between nodes depicts confidence score, with a thicker line representing a higher score. HLA-DRB3 was not recognized by the software and thus excluded from the interaction network. (F) Number of differentially expressed genes from pairwise comparisons of latent cells to all other samples. Differential target expression between samples was performed with a p-value cutoff at < 0.05. (G) Venn diagram displaying the overlay of differential expression hits from comparisons of latently-infected cells with all other samples using a p-value cutoff of p < 0.05. (H) Bar graphs depict the log_2_ fold change of the indicated targets between latent cells and all other samples. Error bars show standard deviation (SD).

Expression data were then adjusted for confounding donor effects for downstream analyses. Subsequently, we examined changes in regulatory and signaling pathways in latent cells based on undirected and directed global significance scores (Figure 3B). The ‘Antigen Processing’, ‘Adhesion’, ‘Pathogen Defense’, and ‘Interleukins’ pathways were the most significantly modulated pathways in the latent compartment. The upregulation of ‘Interleukins’ and ‘Pathogen Defense’ in latently-infected cells likely reflects intracellular signaling cascades that antagonize productive infection. The suppression of ‘Antigen Processing’ in latent cells indicates the ability to evade host immunity, a pro-survival effect that could enforce viral persistence. This observation is in line with previous studies reporting that HIV actively subverts antigen presentation pathways^30,31^.

Next, overall target expression in the sorted samples was assessed by differential expression analysis (DE, Figure 3C). Comparison of uninfected and latently-infected cells revealed 16 differentially expressed targets (adjusted p-value < 0.1), two of which were identified at the protein level: HLA-DRA protein was downregulated, and NT5E (CD73) protein was upregulated on the surface of latent cells. DE analysis of productively- and latently-infected cells resulted in 36 significant hits (adjusted p-value < 0.1), including cell-surface CD4 protein and CD73 protein. CD4 protein was significantly downregulated in productively-infected cells (as expected in the setting of viral expression), and CD73 protein was again upregulated in latent cells.

Overlapping both individual DE analyses revealed 27 targets that were uniquely modulated in latent infection (p-value < 0.05, Figure 3D) with respect to both productively-infected and uninfected cells. These targets comprised CD2 mRNA, a checkpoint molecule which was reported previously as a surrogate marker for HIV reservoirs^14^, and interestingly, CD73 as only differentially-expressed protein (Table S2). The list further included three mRNAs encoding transcription factors, six encoding cytokines, and 11 encoding surface proteins. Among these, *NFKBIA* (NF-κB Inhibitor Alpha) mRNA, which encodes a master inhibitor of the transcription factor NF-κB^32^, a crucial transcription factor of HIV^33,34^, was significantly upregulated in latent cells. In addition, *CASP1* mRNA was significantly downregulated in latently-infected cells. CASP1 (Caspase-1) is a key regulator of pyroptotic cell death, a highly inflammatory process that has been reported to be a major determinant of HIV pathogenesis and a potent driver of virus-dependent CD4+ T cell depletion^35^.

The relationship between molecular markers of HIV latently-infected cells was then investigated by utilizing the open-source software Cytoscape to generate a detailed protein-protein interaction network (Figure 3E). Interestingly, all factors associated with HIV latency in our analyses were members of a single, established protein interaction complex. The chemotactic factor IL8 (or CXCL8) was found to have the highest number of immediate connections (17 first neighbors), followed by the transcription factor STAT1 (Signal Transducer and Activator Of Transcription 1, 15 first neighbors) and the cytokine CSF2 (Colony Stimulating Factor 2, 15 first neighbors).

### Three key members of the adenosinergic pathway, IL8, CD39, and CD73, are differentially expressed in latent cells

To deepen our characterization of HIV latently-infected cells, we additionally considered non-activated (no exposure to αCD3/αCD28-stimulating beads) and non-virus-exposed cell populations in our NanoString analyses. Inclusion of these comparators more faithfully emulates the *in vivo* landscape in ART-suppressed individuals where the majority of CD4+ T cells are likely in a resting state and have not encountered HIV.

Next, DE analysis was implemented to compare latent cells in a pairwise fashion to all other cellular populations (Figure 3F). The pairwise DE analyses were then overlaid to identify genes that were uniquely regulated in latently-infected cells (Figure 3G). We identified three targets that were significantly modulated with respect to all other comparator groups: IL8 mRNA, CD39 mRNA, and CD73 protein (a key marker revealed in our previous analyses) (Figure 3H). These three factors are known to be mechanistically connected through the adenosine signaling pathway, and our observed induction of IL-8 gene expression in latently-infected cells (Figure 3H) potentially reflects ADORA2B stimulation^36–40^

### CD73 is highly expressed on a subset of PD-1, CD39, CD49d and CD2-positive cells

CD39 and CD73 are surface proteins and thus could serve as convenient indicators of latent reservoir cells. We therefore sought to examine correlations between the surface expression of CD73 and previously reported reservoir markers and enrichment factors, including CD2, CD49d, PD-1, CD39, CD98, CTLA-4, CD20, CD30 and CD32 in blood-derived CD4+ T cells from people living with HIV (PLWH). CD4+ T cells were negatively-selected from fresh blood collected from six PLWH on ART, and reservoir markers were measured by immunofluorescence staining and flow cytometry. Notably, none of the previously identified reservoir markers were enriched in the CD73+ CD4+ T cell compartment, and the frequency of PD-1 and CD39 expression was significantly reduced in CD73+ cells as compared to total CD4+ cell populations (Figure S3 A and B). However, we also identified double-positive cells co-expressing high levels of CD73 and CD49d, CD2, PD-1, or CD39, respectively (Figure S3 C, red arrows), indicating that these markers are not mutually exclusive *in vivo* in the setting of treated HIV infection. Expression levels of CD98, CTLA-4, CD20, CD30 and CD32 were extremely low in both total CD4+ T and CD73+ CD4+ T cells (Figure S3, B-D).

Based on unsupervised analyses including principal component analysis (PCA), expression signatures were dominated by *in vitro* stimulation (Figure S4 A and B). We therefore investigated T cell activation as a potential determinant of HIV latency by examining the expression levels of established T cell surface markers CD45, CD45RO, and CD4 (Figure S4 C), as well as panel-specific activation markers (Figure S4 D and E). No significant associations were discovered between the expression levels of these activation markers and latent infection, suggesting that viral transcriptional fates are not dictated by cellular activation states.

### Hypoxia promotes CD73 expression and HIV latency in primary CD4+ T cells

To better understand the relevance of CD73 in the context of HIV infection and particularly in the establishment of the latent reservoir, we next focused on the regulation of CD73 expression^41,42^. Importantly, at least one hypoxia-response element (HRE) has been previously identified in the CD73 promoter region^43^ allowing for direct binding of hypoxia-inducible factors (HIFs). In this context, HIF-1 was demonstrated to control CD73 expression, with CD73 typically being upregulated under hypoxic conditions^44^. We therefore hypothesized that our discovery of CD73 upregulation in latent cells suggested that hypoxia may be the underlying cause of both increased CD73 expression and HIV latency.

We first confirmed our NanoString data using *in vitro* infection with HIV_DFII_ followed by immunofluorescence staining and flow cytometry, again observing a significant enrichment of CD73+ cells among latently-infected cells (Figure 4A and B). We then mimicked hypoxic conditions in our culture settings via administration of dimethyloxalylglycine (DMOG), followed by infection with HIV_DFII_ and flow cytometry (Figure 4C).

**Figure 4:**
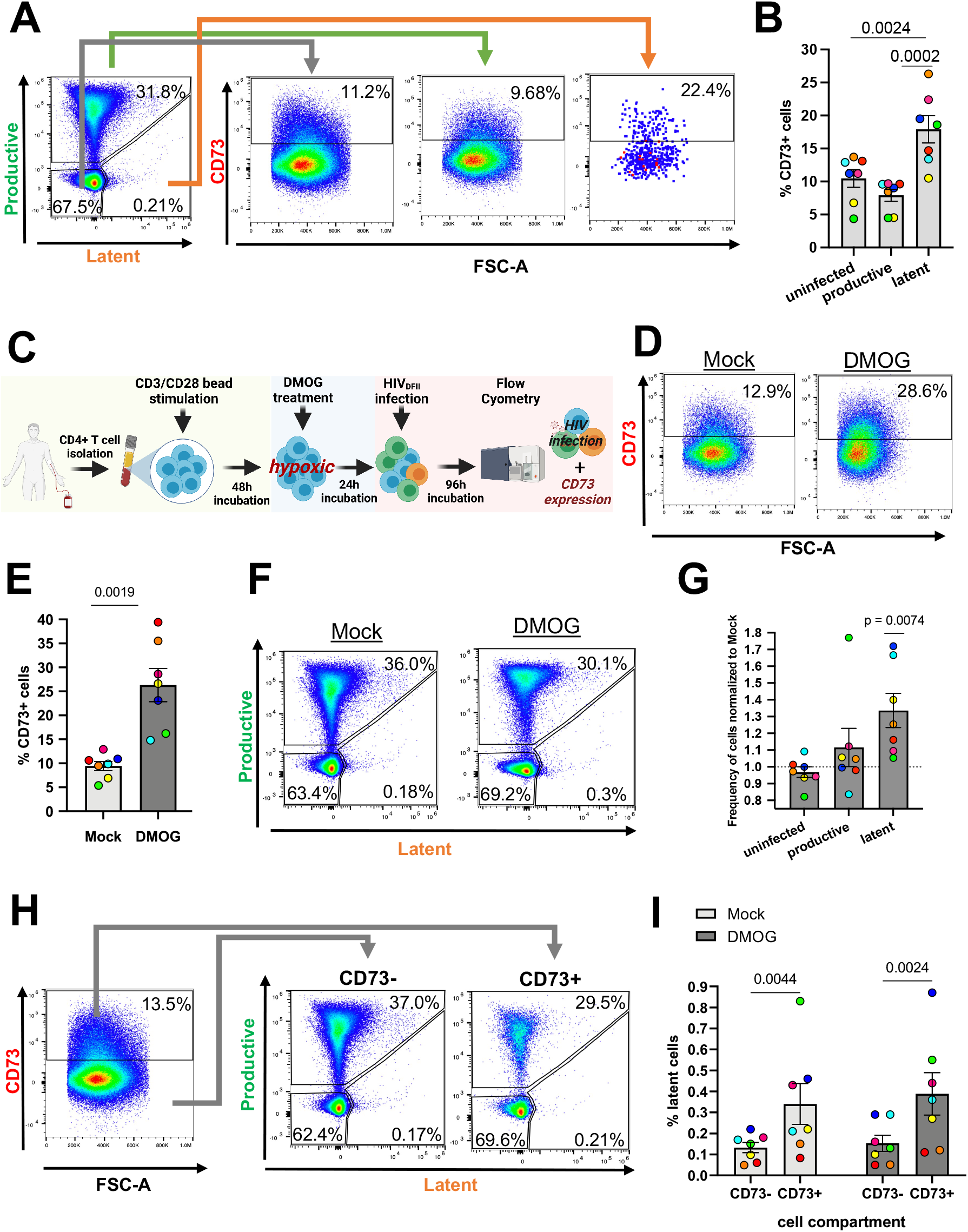
Hypoxic conditions increase the frequency of CD4+ CD73+ T cells and facilitate latent infection. (A) CD4+ T cells from healthy individuals were infected with HIV_DFII_ and analyzed by immunofluorescence staining and flow cytometry. Scatter plots show the gating strategy to analyze CD73 expression in HIV_DFII_-infected cells from a single, representative donor. (B) Shown is the frequency of CD73+ cells in the respective HIV_DFII_-infected cell population summarized from 7 donors. P-values displayed above bars were generated by performing a one-way ANOVA with Tukey’s multiple comparisons test. (C) Experimental workflow for HIV_DFII_ infection under hypoxic conditions. (D) Representative flow plots of CD73 immunofluorescence staining in mock or DMOG treated CD4+ T cells and (E) average frequency of CD73+ CD4+ T cells across 7 donors. The p-value was generated by performing a two-tailed paired t-test. (F) Representative flow plots, showing HIV_DFII_ infection profiles 4 days post HIV_DFII_ exposure upon mock or DMOG treatment. (G) The frequency of DMOG-treated, HIV_DFII_-infected cells was normalized to the respective mock population within each donor to account for donor variability. P-values were generated by performing a two-way ANOVA with Fisher’s Least Significant Difference (LSD) test. (H) Gating strategy used to analyze HIV_DFII_ infection profiles in the CD4+ CD73+/− T cell compartments. Data shown are from a single, mock-treated donor and are representative for all seven donors. (I) The average frequency of latently-infected cells was assessed in CD4+ CD73+/− T cells. Colored dots indicate individual donors. P-values were generated by performing a two-way ANOVA with Fisher’s LSD test. P-value threshold for significance was at p < 0.05. Error bars show SEM.

DMOG treatment substantially altered CD73 expression in CD4+ T cells and led to a 2.5-fold increase of CD73+ cells compared to mock treatment (Figure 4D and E). Similar levels of CD73 induction were observed in cells incubated at 1% oxygen, indicating that DMOG treatment accurately recapitulates the CD73 phenotype in CD4+ T cells (Figure S5A). Notably, while CD73 expression was generally increased upon induction of hypoxic responses (compare Figure 4B and Figure S5B), we observed a pronounced enrichment of CD73+ cells specifically in the latent compartment with up to 60% of latent cells being CD73 positive (Figure S5B). This was also reflected in CD73 expression per cell measured by mean fluorescence intensities (MFIs), which showed an overall increased CD73 expression in latent cells in both mock and DMOG-treated samples (Figure S5C). Latent infection became significantly more abundant with a 33.6% increase upon DMOG treatment, while the frequency of uninfected or productive cells did not change significantly between culture conditions (Figure 4F and G).

Lastly, we examined whether CD73+ cells were enriched for latent virus (Figure 4H). We measured a considerably higher rate of latently-infected cells in CD73+ compared to CD73-cells with an average 3-fold enrichment of latent infection within the CD73+ T cell compartment (Figure 4I). Considering the average frequency of CD73+ cells among CD4+ T cells (∼10%, Figure 4E), these data suggest that approximately 25% of the overall, peripheral CD4+ T cell HIV reservoir may reside in CD73+ cells. Interestingly, the enrichment of latent cells in CD4+ CD73+ T cells did not significantly differ between mock- and DMOG-treated samples (Figure 4I), indicating that CD73+ cells may possess specific features that favor the establishment and/or maintenance of latent infection that are not further modulated by the induction of hypoxic responses.

### CD73+ cells exhibit distinct immunoregulation and tyrosine kinase signaling

The role of CD73 in the context of oncogenesis, tumor progression and survival is well described^38^. However, relatively little is known about the relevance and function of CD73 in human CD4+ T cells. We thus characterized the transcriptome of blood-derived, primary CD73+ and CD73-CD4+ T cells by RNA sequencing. To address this aim, CD73-/+ cells were isolated from primary CD4+ T cells by FACS, revealing highly variable frequencies of CD73+ CD4+ T cells across donors, ranging from ∼3% to ∼23% (Figure S6A and B). Sorted cells were then subjected to RNA sequencing and DE analysis, accounting and adjusting for donor effects (Figure S6C and D). CD73- and CD73+ CD4+ T cells exhibited distinct transcriptional profiles and a clear clustering of the sorted populations (Figure 5A). Expectedly, the CD73 (*NT5E*) exhibited highly significant divergence in expression levels between groups (Figure 5B). An additional 145 genes were differentially expressed, with 111 upregulated and 34 downregulated genes in CD73+ cells (Figure 5B). *CR1, ADAM23, ABCB1 and AUTS2* were among the top genes upregulated in CD73+ cells (Figure 5B, yellow dots)*. CLEC17A* exhibited the highest fold change (> 5-fold), followed by *CD73, LINC02397* and *MACROD2* (> 4-fold) (Figure 5B, red dots). Multiple genes were markedly downregulated in CD73+ cells; however, only two of them reached the highest level of statistical significance (adjusted p < 0.01): *FCER1A* and *NPR3*. A gene set enrichment analysis yielded a multitude of significantly modulated pathways differentiating CD73+ and CD73-cells including ‘immune responses’, ‘complement activation’ and ‘receptor-mediated signaling cascades’ (Figure 5C), indicating diverging immunoregulatory programs. In addition, ‘positive regulation of angiogenesis’, ‘regulation of apoptotic processes’ and ‘tyrosine kinase signaling cascades’ were among the 40 most significantly modulated pathways.

**Figure 5:**
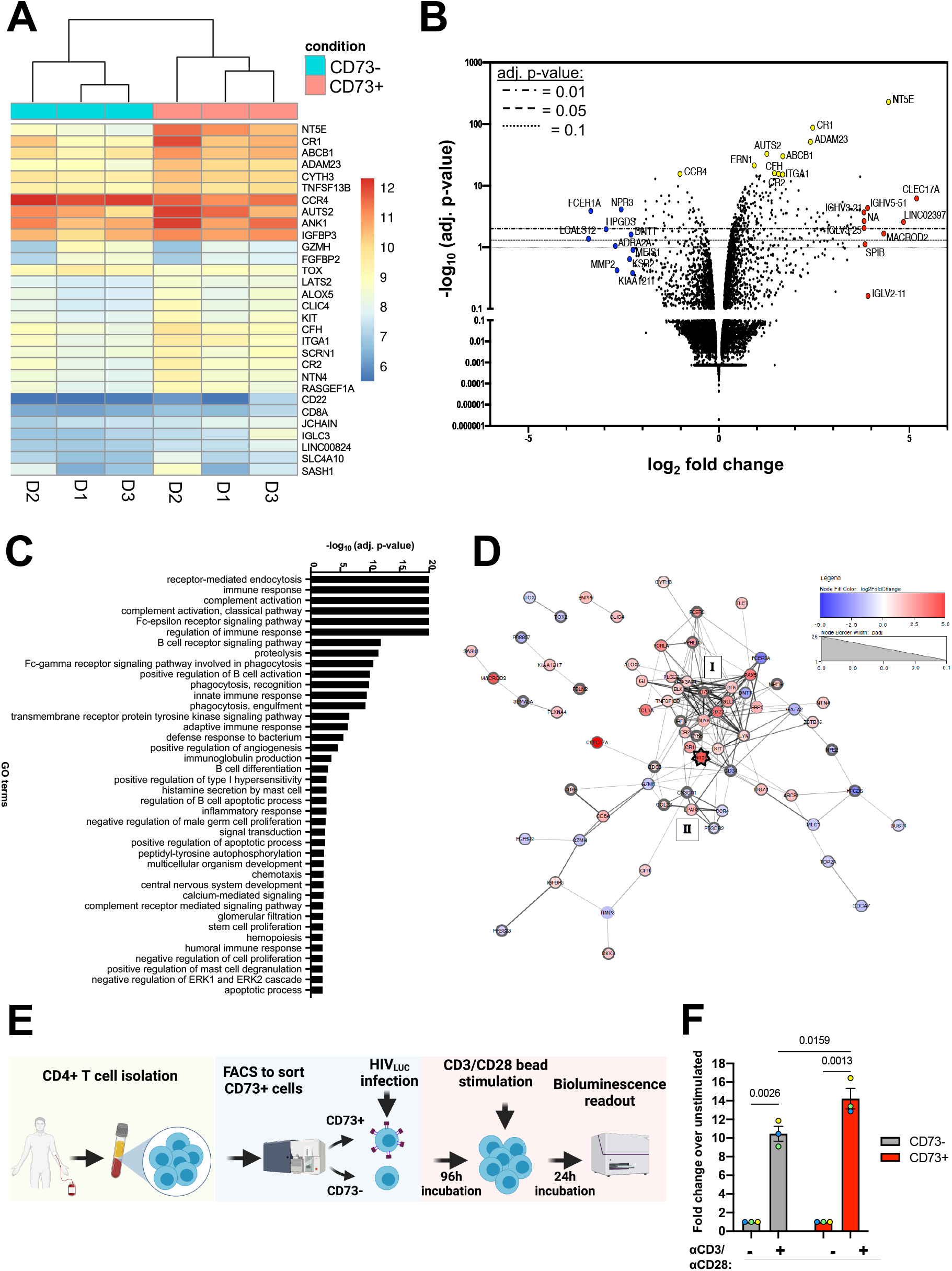
CD73+ cells exhibit distinct immunoregulatory features and support HIV reactivation. (A) Clustering heatmap visualizing the expression profile of the top 30 differentially-expressed genes sorted by their adjusted p-value. Data were adjusted for donor effects. (B) Global transcriptional changes between CD73-and CD73+ cells are visualized in the volcano plot. Each data point represents a single gene. The log_2_ fold change of the normalized mean hit counts of each gene are plotted against -log_10_ of its adj. p-value (using the Benjamini-Hochberg method). Horizontal lines indicate adj. p-value thresholds. The top 10 most significantly changed (yellow), most upregulated (red) and most downregulated (blue) genes are highlighted. (C) Differentially expressed genes were grouped by their gene ontology (GO) and the enrichment of GO terms was tested using Fisher exact tests. The top 40 GO terms with an adj. p-value < 0.05 are displayed. (D) The protein-protein interaction network of genes differentially expressed between CD73- and CD73+ CD4+ T cells was generated in Cytoscape as described previously. 33/145 hits were not recognized by the STRING database and thus excluded from the interaction network. Among the remaining 112 genes, 43 genes did not have an interaction partner and are not displayed. Log_2_ fold changes of genes are indicated by node pseudo color and the adj. p-value by node border width. *CD73* (*NT5E*) is highlighted by a star shape. Protein clusters of interest are labeled. (E) The schematic represents the experimental workflow to test reactivation of integrated provirus in sorted CD73- and CD73+ CD4+ T cells. (F) Bars display the average fold change over unstimulated cells from three donors. Colored dots indicate individual donors. P-values above bars were generated by performing a two-way ANOVA with Fisher’s LSD. A p-value at p < 0.05 was considered significant. Error bars show SEM.

Next, all differentially expressed genes were uploaded into Cytoscape and the resulting protein-protein interaction network was then overlaid with log_2_ fold changes and adjusted p-values of each gene, visualizing directionality and significance of interacting hits (Figure 5D). The analysis revealed five distinct interaction networks, with one network containing the majority of hits. This approach revealed biological programs associated with the CD73+ CD4+ T cell compartment. Within this network, a tight cluster (I) of cell surface receptors was apparent (*CD22, CD79A, CR1, CR2, KIT, FCER1A, CD34*), accompanied by several signaling factors (*LYN, BLNK, BLK, BTK, PIK3AP1, TCL1A)* and transcription factors (*IRF8, EBF1, PAX5*). Another cluster (II) comprised downregulation of cell surface proteins *CXCR3*, *CCR4* and *PTGDR2*. Importantly, the network also contained two significantly downregulated genes, transcription factor *GATA2*, and DNA topoisomerase *TOP2A*, which are known to support active HIV transcription and replication^45,46^.

### CD73+ CD4+ T cells harbor an inducible HIV reservoir

The intactness of integrated proviruses and the inducibility of viral gene expression are key attributes of the latent HIV reservoir. We therefore sought to investigate the capacity of CD73+ T cells to harbor reactivatable latent virus. To this end, we adapted a primary *in vitro* HIV infection model^47^ and infected FACS-sorted CD73- and CD73+ CD4+ T cells with the reporter virus HIV_Luc_ (Figure 5E). After a resting period of 5 days, we stimulated cells with αCD3/αCD28 beads and observed a pronounced increase of viral transcription in both CD73- and CD73+ CD4+ T cells as compared to unstimulated cells, reflected by a 10-fold and 14-fold increase in LTR-driven luciferase activity, respectively (Figure 5F). Induction of viral transcriptional activity was significantly higher in stimulated CD73+ T cells as compared to CD73-T cells. These results demonstrate that the CD73+ CD4+ T cell compartment can harbor an inducible latent reservoir *in vitro*, indicating the potential of this compartment to contribute to spreading infection in people PLWH upon treatment cessation.

### Blocking the adenosine receptor A2AR or CD73 promotes HIV latency reversal

Bearing in mind the enzymatic function of CD73, we hypothesized that extracellular adenosine produced by CD73 could be mechanistically involved in the establishment and/or maintenance of HIV latency. To test this, we investigated the adenosine signaling cascades downstream of CD73 in the well-established J-Lat cell line models of latency^48,49^, exploring how modulation of adenosine receptors by small molecule drugs affects HIV transcriptional activity.

First, CD73 surface expression was measured in J-Lat clones, 5A8, 6.3, 11.1, and A72 as well as in the parental JurkatE6 cell line (Figure 6A). We found that between 10-20% of JurkatE6, J-Lat 6.3, 11.1, and A72 cells expressed CD73 on the surface, whereas about 80% of J-Lat 5A8 cells were CD73 positive. In the next experiment we sought to test whether an inhibition of adenosine signaling will impact HIV transcription in J-Lat cells. To that aim, J-Lat 5A8 cells were pretreated with the adenosine receptor antagonist SCH-58261 (SCH), followed by viral reactivation with PMA/I, a strong mitogen and latency reversing agent. Notably, pretreatment with SCH significantly promoted latency reversal and resulted in an evident dose response with a 2-fold increase in the frequency of GFP+ cells at the highest dose (Figure 6B). At the same time, SCH pretreatment did not compromise cell viability at any dose compared to cells treated with PMA/I only (Figure 6B).

**Figure 6:**
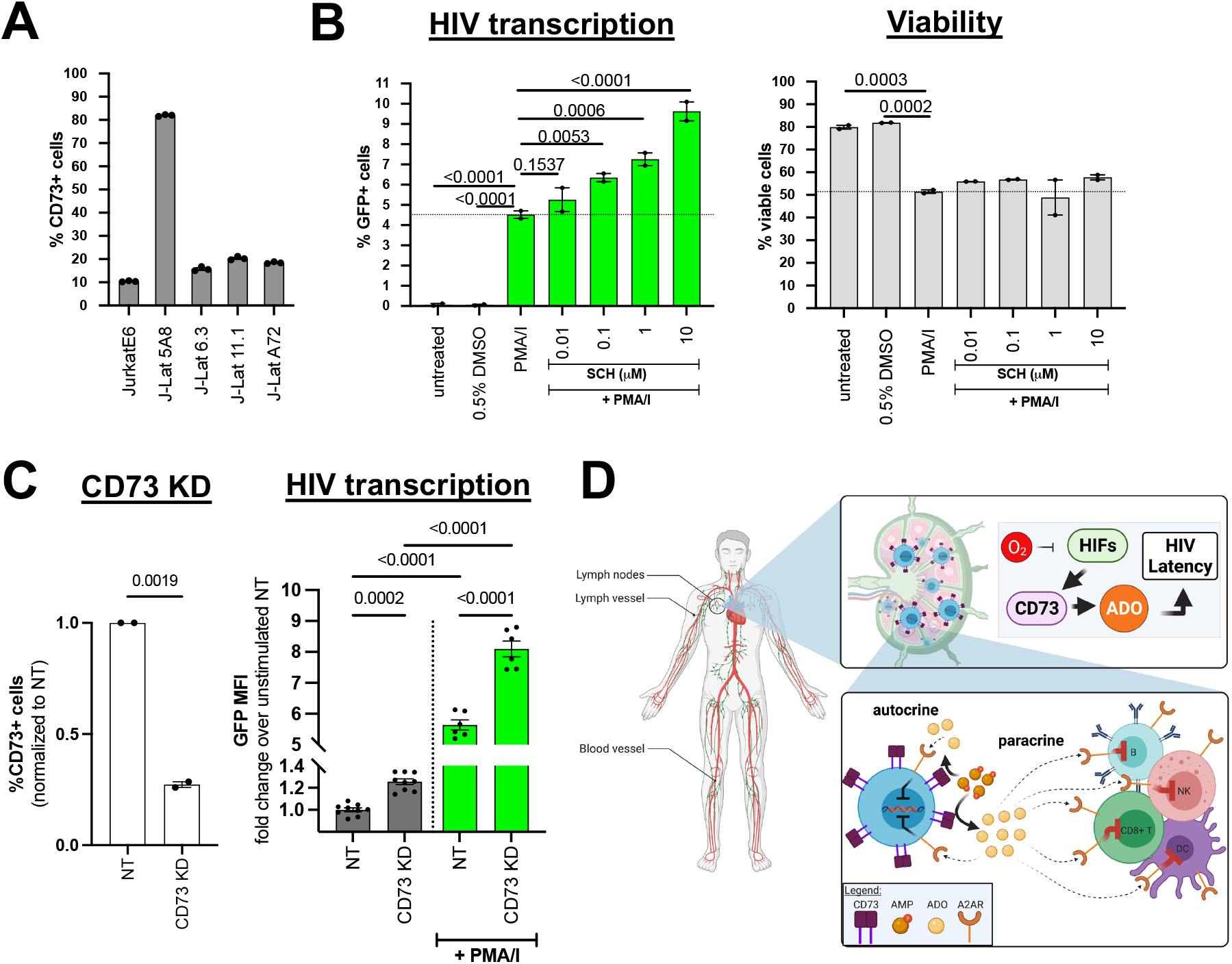
Blockade of CD73 and adenosine receptor A2AR facilitates HIV latency reversal. (A) Average frequency of live, single CD73+ cells in the different T cell lines measured via flow cytometry. (B) J-Lat 5A8 cells were treated with A2AR antagonist SCH-58261 (SCH) at indicated, increasing doses for 1h, followed by treatment with PMA/I. PMA/I treatment alone served as positive control, 0.5% DMSO treatment as mock control. Cell viability based on FSC-A and SSC-A and GFP expression as a reporter of HIV transcriptional activity was assessed 7h post treatment using flow cytometry. Samples were measured in duplicates. P-values were generated by performing a one-way ANOVA with Fisher’s LSD. p < 0.05 was considered significant. Data represent the mean and error bars the SEM. (C) J-Lat A72 cells were transduced with lentiviral vectors containing dCas9-KRAB sgRNAs targeting CD73 or a non-targeting control (NT). CD73 KD was assessed via immunofluorescence staining of CD73 as measured by flow cytometry. Data were normalized to NT. Reactivation of viral transcription reflected by GFP expression was assessed by flow cytometry of A72 KD cells in presence and absence of PMA/I. Significance was determined by one-way ANOVA, comparing samples with NT. (D) HCA-model of HIV persistence. Under long-term ART, HIV persists in vivo preferentially in anatomical sanctuary sites like lymph nodes. Oxygen levels are known to be significantly lower in lymphoid tissues (0.5-4.5%) compared to the periphery (about 13%). Low oxygen supply or hypoxia lead to stabilization of hypoxia inducible factors (HIFs) which control the expression of CD73. Upregulation of CD73 leads to the accumulation of adenosine (ADO) in the extracellular space. These factors favor the establishment of viral reservoirs and promote HIV persistence, silencing the provirus through autocrine signaling and inhibiting host antiviral immune responses by paracrine signaling mechanisms. T cells are shown in blue, CD73 in purple.

Finally, we sought to determine if downregulation of CD73 itself affects HIV transcription and latency reversal. To pursue this objective, we performed CD73 knockdown (KD) via CRISPR interference (CRISPRi) in the A72 J-Lat clone. We first introduced the CRISPRi effector protein dCas9-KRAB by lentiviral transduction, generating the cell line J-Lat A72 CRISPRi. We then utilized another lentiviral vector to express single guide RNAs (sgRNAs) targeting CD73 or a non-targeting (NT) negative control, respectively. We tested KD efficiency by immunofluorescence staining of surface CD73, finding a significant downregulation of CD73 compared to non-targeting controls (Figure 6C). CD73 KD resulted in significantly lower GFP expression levels in both stimulated and non-stimulated conditions (Figure 6C), indicating that CD73 exerts direct effects on HIV transcription and latency.

### A comprehensive model of CD73-dependent latency

Based on our data obtained in *in vitro* infection models, we devised a model that incorporates hypoxia, CD73, and adenosine signaling, linking these factors with HIV transcriptional regulation and persistence. Our model predicts that the hypoxia-CD73-adenosine (HCA) axis plays a vital role with CD73 as a central player in the maintenance of latent HIV infection (Figure 6D): As is known, HIV preferentially persists in lymphatic tissues^50^, which exhibit low oxygen levels and thus high levels of active HIFs. We surmise that the resulting upregulation of CD73 expression leads to adenosine-rich, immunosuppressive microenvironments, which promote the establishment and maintenance of latent reservoirs. Accumulation of extracellular adenosine thereby creates optimal conditions for HIV persistence by suppressing HIV host dependency factors through autocrine signaling cascades, while impairing effective host immune responses through paracrine signaling mechanisms. Therefore, viral transcription is repressed, and immune escape of HIV-infected cells is facilitated, altogether promoting viral quiescence and survival of the latent reservoir.

### Detection of HIV markers and CD73 *in vivo* in tissues from ART-suppressed HIV-infected individuals

To evaluate the clinical relevance of our findings and to test the HCA model *in vivo*, we applied a comprehensive tissue imaging pipeline that enables simultaneous detection of integrated HIV-DNA, viral mRNA, and HIV protein, as well as lineage markers, followed by quantitative image analyses^51–53^. We concomitantly detected CD73 expression and HIV markers in lymph node tissues (peripheral and inguinal lymph nodes) obtained from HIV-infected individuals on suppressive ART (ART suppr.), as well as in viremic untreated individuals, and uninfected controls (Figure 7A). CD3 staining was used as a lineage marker for T cells and DAPI was included as a counter staining of nuclear DNA. CD73 signals were found across all samples, while HIV detection expectedly differed greatly between ART-suppressed and viremic individuals and was absent in uninfected control samples (Figure 7A and B). Quantification of the viral reservoir in viremic individuals indicated that at least 40% of CD3+ cells harbored integrated DNA and most of these cells expressed viral RNA and HIV-p24 (Figure 7B, viremic). In contrast, in ART-suppressed individuals, 0.017% of CD3+ cells contained viral DNA, half of which expressed viral RNA and 10% of which expressed viral proteins (Figure 7B, ART-suppr.).

**Figure 7:**
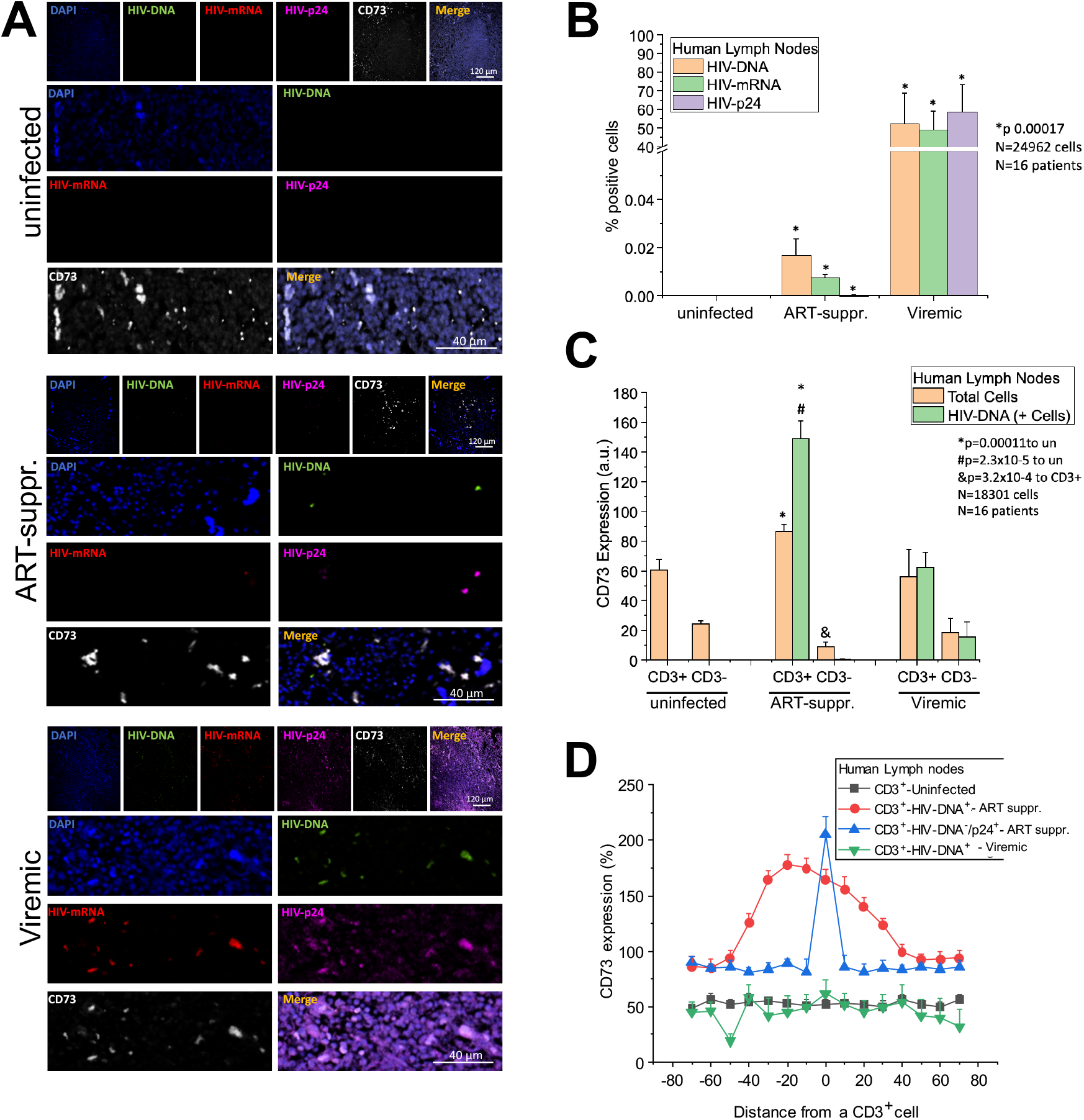
*In situ* imaging of lymph nodes from PLWH reveals associations between CD73 protein expression and HIV DNA specifically in ART-suppressed individuals. (A) Panels show representative confocal microscopy images for CD73 and HIV marker staining of lymph node tissue obtained from uninfected, ART-suppressed (suppr.), and viremic individuals. (B) Automated image analysis of tissue sections stained for HIV DNA, mRNA and p24 protein in uninfected (n = 5), ART-suppressed (n = 6), and viremic individuals (n = 5). Bar graphs show the average percentage of cells that exhibit a positive signal for the respective target. (C) Quantitative CD73 expression analysis in tissue samples from uninfected (n = 5), ART-suppressed (n = 6) and viremic individuals (n = 5). Identification of the T cell compartment (CD3+ cells) and detection of HIV reservoir cells (HIV-DNA+) was followed by quantification of CD73 signals in the indicated 4 cellular subsets (CD3+ HIV-DNA-, CD3+ HIV-DNA+, CD3-HIV-DNA-, CD3 HIV-DNA+). ANOVA was used to compare groups. Relevant statistics are shown in bar graphs. (D) Quantification of CD73 expression and distribution in and around viral reservoirs in µM. Lineplot analysis was performed as described in the Methods section and the schematic shown in Figure S7. 40-98 3D reconstructed images were analyzed per condition from 16 independent participant samples. CD73 intensity values were normalized to mean intensities in CD3+ HIV-DNA-/p24+-ART suppr. samples.

### CD73 expression correlates with HIV persistence in lymph nodes from ART-suppressed individuals

Next, we determined the association between HIV infection and CD73 expression, specifically in the CD3+/CD3-cell compartment (Figure 7C). CD73 expression levels in CD3+ cells were significantly higher in ART-suppressed samples than in uninfected samples. In addition, HIV DNA+ CD3+ cells in ART-suppressed individuals exhibited a significant, 1.6*-*fold higher CD73 expression compared to the overall CD3+ population. This difference in CD73 expression levels was not observed in viremic individuals and suggested that CD73 expression may offer a survival advantage for HIV-infected cells in the setting of ART. We further hypothesized that viral reservoirs would not only reside in individual CD73+ cells, but rather in CD73-rich, HIV persistence-promoting microenvironments, due to local hypoxia and paracrine adenosine signaling effects. To explore this hypothesis, we performed imaging analysis by measuring the expression of CD73 in CD3+/HIV-DNA+ cells as well as neighboring uninfected cells surrounding viral reservoirs (Figure 7D and Figure S7). Zero represents foci of viral infection, and increasing or decreasing distances correspond to neighboring uninfected cells (Figure 7D, up to 80 µm). Most localized expression of CD73 was observed in and around viral reservoirs (CD3+HIV-DNA) in ART-suppressed individuals. In addition, we observed that CD3+ cells that were negative for HIV-DNA but HIV-p24 positive (HIV-DNA^-^/p24^+^), perhaps due to the uptake of viral particles, also had increased levels of CD73. In contrast, evenly-distributed expression of CD73 was observed in uninfected and viremic lymph node tissues, confirming that the spatial association between CD73 expression and viral reservoirs is specific to ART-suppressed HIV infection (Figure 7D, black and green lines, respectively). Our data indicate that HIV DNA+ CD3+ cells in lymph nodes from ART-suppressed individuals reside in areas with generally high CD73 expression, reinforcing that CD73 via adenosine production contributes to HIV persistence (Figure 7D).

## DISCUSSION

The latent HIV reservoir is considered to be the main obstacle to achieving viral eradication or indefinite disease remission in the absence of ART. It is generally believed that integrated provirus persists in specific cellular compartments and tissue sanctuaries which enable both long-term survival and spontaneous reactivation of viral replication even after decades of successful ART. The exact identity of the HIV reservoir however remains elusive. The goal of this study was to thoroughly characterize primary HIV latently-infected cells using comprehensive and complementary systems approaches. Our work focused on CD4+ T cells, the major target of HIV infection and an important potential source of viral rebound upon ART interruption.

The core finding of this work was the identification of elevated cell-surface expression of CD73 as a signature characteristic of HIV latently-infected CD4+ T cells – an observation that, to the best of our knowledge, has not been described before. Using different *in vitro* models of HIV latency, we found indications that CD73 is not merely a passive marker of reservoir cells, and regulation of CD73 expression as well as its enzymatic function is mechanistically linked to HIV persistence.

Our NanoString analyses also revealed significant changes in the expression of CD39 and IL8 within the latent compartment, which are mechanistically connected to CD73. In particular, the two ectonucleotidases CD39 and CD73 and their orchestrated functionality in purinergic signaling are well described in the literature^38–40^. In addition, IL8 expression has been shown to be stimulated by adenosine signaling, and is thus directly linked to the enzymatic function of CD73^37^. The distinctive expression of these three factors in latent cells suggests that CD73 and the adenosinergic pathway play a pivotal role in HIV latency and the establishment and/or maintenance of viral reservoirs.

Our initial *in vitro* infection model using HIV_DFII_ focused on a rather narrow time window of HIV latency, recapitulating the establishment and early persistence of viral reservoirs within a few days after infection. In the setting of HIV infection *in vivo*, HIV persists even after decades of fully suppressive ART. Linking the two observations, our *in situ* imaging of lymph node tissues demonstrated, for the first time, that a significant association between CD73 expression in T cells and HIV-DNA specifically in ART-suppressed individuals is maintained over extended periods of time. No spatial associations between CD73 and HIV markers were observed in tissues from viremic, untreated individuals. We surmise that in the absence of treatment, when HIV is not subject to pharmacologic selection pressure, active viral replication overrides CD73 effects and leads to rapid turnover of infected cells. In the setting of long-term ART, HIV-infected cells are subject to immune selection pressure, leading to an enrichment in CD73+ infected cells due to their immunosuppressive nature and their increased likelihood of harboring transcriptionally inactive virus. This hypothesis is further reinforced by our transcriptomic profiling of blood-derived CD73+ and CD73-CD4+ T cell subsets which revealed a spectrum of biological programs in the CD73 compartment that are expected to promote viral quiescence, immune evasion, and cell survivorship.

A key finding of our study is that CD73 is significantly upregulated under hypoxic conditions in the CD4+ T cell compartment. This result is important for two reasons: 1) Oxygen tension levels are highly variable throughout the body, ranging from 110 mmHg in well oxygenated lung alveoli^54^, down to 55 – 60 mmHg^55^ in the gastrointestinal tract, and 3 – 35 mmHg in lymphoid organs^56,57^. As a consequence, immune cells including CD4+ T cells encounter and operate at varying oxygen concentrations as they traffic through the body^57^. Lymphoid organs in particular represent crucial viral sanctuaries and as these are characterized by low oxygen levels, our data suggest a causal link between hypoxia, CD73 expression, and HIV persistence. 2) Hitherto, the role of oxygen levels and hypoxia in HIV infection remained intangible and somewhat obscure. In 2009 Charles et al.^58^ reported decreased HIV-1 RNA levels at 3% oxygen, while S. Deshmane and colleagues showed increased HIV transcription mediated by the interaction between HIV accessory protein Vpr and HIF-1α^59,60^. Moreover, HIV-1 replication was shown to be promoted by HIF-1α which in turn was stabilized by reactive oxygen species^61^. Very recently it was demonstrated that hypoxia can promote HIV latency with HIF-2α as a direct inhibitor of viral transcription^62^. Our data now provide new insights into the connection between HIV latency and hypoxia, with CD73 emerging as key mediator bridging hypoxia and viral transcription.

As outlined above, our data suggest mechanistic involvement of CD73 in HIV persistence via its enzymatic function in the adenosine signaling cascade. Adenosine signaling leads to a suppression of cellular transcription factors that are critical for active viral transcription^63,64^ and may create an ideal immunological niche for infected cells to evade host immune clearance due to an adenosine-mediated immunosuppressive microenvironment. Pharmacological blockade of A2AR in our latency model facilitated HIV latency reversal without impairing cell viability. Thus, our data demonstrate the potential of targeting the adenosinergic system as a therapeutic approach in the context of HIV infection and provides first evidence that the enzymatic activity of CD73 is directly involved in HIV persistence. We would like to point out that hypoxia can regulate the expression of numerous genes^65^, including, but not limited to CD39, the adenosine receptor ADORA2B, the equilibrium nucleoside transporter ENT-1 and adenosine kinases (ADK)^66^, all of which could promote adenosine signaling independent of, or in concert with CD73. Additional studies will therefore be necessary to determine how these diverse factors link hypoxia-mediated signaling to HIV persistence.

It is important to note that in recent years increasing circumstantial evidence has been gathered, independently suggesting a critical role for either hypoxia^58–62^, CD73 expression^67–69^ or adenosine signaling^70–74^ in HIV infection. Our findings reported here are complemented and validated by a recent translational study by Seddiki *et al.* demonstrating that HIV-infected CD73+ memory CD4+ T cells contribute significantly to the very long-lived HIV proviral DNA reservoir in treated subjects^75^. Our data now reveal a direct, causal connection that links the hypoxic regulation and adenosine-producing enzymatic activity of CD73 to the establishment and persistence of viral reservoirs.

Finally, the CD73-adenosine axis has received a great deal of attention in the context of cancer biology^76–78^ and the adenosinergic system has emerged as a promising new drug target in oncology^37,38,79–84^. Our data suggest that, like solid tumor cells, latent HIV reservoirs hijack the CD73-adenosine axis to subvert innate and adaptive immune responses, enhancing the survivorship of the infected cell as a persistence mechanism during ART. Therefore, CD73- and adenosine-focused anti-cancer therapeutics should be actively explored as foundations of novel HIV cure approaches and host-directed therapies.

### Limitations of the study

There are limitations to both our in vivo models and our in vivo profiling work that should be considered. Firstly, our in vitro HIV latency models are short-term infection models, which were mainly geared at identifying host factors associated with the establishment of viral transcriptional latency in newly-infected cells. These models do not capture the critical aspects of immunological selection that occur in vivo over the course of years or even decades of suppressive ART in PLWH; long-term selection pressures in vivo may result in enrichment of host factors on the surface of latently-infected cells that do not appear in our in vitro profiling experiments. In regards to our in vivo profiling of lymphoid tissues, analyses of a much larger collection of tissue samples (from a broader array of anatomic sites associated with HIV persistence) will be required to fully appreciate the association between CD73 expression and the HIV reservoir. Moreover, our studies here have not considered HIV persistence outside of the lymphoid compartment; as there is evidence that myeloid lineage cells (e.g. monocyte-derived macrophages) harbor HIV, the relevance of adenosine signaling to HIV infection should be evaluated in these cell types as well. Lastly, although our in vitro experiments provide mechanistic connections between adenosine signaling and HIV persistence, much additional work remains to be done to decipher the relationship between adenosine signaling, HIV latency, and antiviral immunity in vivo. These future studies will be critical in guiding the development of novel HIV cure strategies targeting the adenosine pathway.

## MATERIALS AND METHODS

### Construction of next generation HIV_DFII_ reporter plasmid

The 1^st^ generation dual reporter virus construct R7GEmC^26^ (kindly provided by Dr. Eric Verdin) was adapted by ligation-based molecular cloning. Briefly, R7GEmC was linearized by enzymatic digest using FseI and AscI (New England Biolabs, Cat. #R0588S and Cat. #R0558S) in order to excise the mCherry open reading frame. Then, mKO2 was PCR amplified from the template mKO2-N1 (Addgene, Cat. #4625) using primers containing matching restriction sites. Subsequently, the lentiviral vector was dephosphorylated using Quick CIP (New England Biolabs, Cat. #M0525S), purified and subjected to ligation with the digested and gel-purified PCR product utilizing T4 DNA ligase (New England Biolabs, Cat. # M0202S).

### Plasmid amplification and preparation

Circularized plasmid DNA was transformed by heat-shock in chemically competent Stbl-3 E. coli (ThermoFisher Scientific, Cat. #C737303) for subsequent antibiotics selection and Sanger sequencing (Elim Biopharm) of positive clones.

Transformed Stbl-3 cells were grown in 2-5 ml of lysogeny broth with 0.1 mg/ml Ampicillin (LB Amp) for 16-24h or 4-8 hours prior to inoculation of large-volume flask with 100-300 ml LB Amp for overnight growth. 16 hours later, cells were pelleted by centrifugation for 15 min at >3000 g and subjected to plasmid preparation using the NucleoSpin Plasmid, Mini kit for plasmid DNA (Macherey-Nagel, Cat. # 740588) or Plasmid Plus Maxi Kits (QIAGEN, Cat. #12963) following the manufacturer’s protocol. Concentrations of isolated plasmid DNA was spectrophotometrically determined using a NanoDrop 1000 (ThermoFisher, Scientific) and DNA aliquots were stored at 4°C.

### Cell lines and cell culture

HEK293T were obtained from ATCC and were cultured in DMEM (ThermoFisher, Scientific, Cat. # 11965-118) supplemented with 10% FBS (Corning, Inc., Cat. #35-010-CV) and 10% Penicillin/1% Streptomycin (Fisher Scientific, Cat. #11548876) (DMEM complete = DMEM+/+) at 37°C, 5% CO_2_ unless stated otherwise. J-Lat 5A8 cells were kindly provided by Warner C. Greene (Gladstone Institutes) and J-Lat A72 cells were obtained from the NIH HIV Reagent Program (Cat. # ARP-9856). All other J-Lat clones were a gift from Eric Verdin (Buck Institute). Jurkat E6-1 cells were obtained from ATCC. All suspension cells were cultured in RPMI 1640 (ThermoFisher, Scientific, Cat. #11875-119) supplemented with 10% FBS and 10% Penicillin/1% Streptomycin (RPMI complete = RPMI+/+) at 37°C, 5% CO_2_ unless stated otherwise.

### Virus production

HIV-1 viruses were generated by transfection of proviral DNA into HEK293T cells via polyethylenimine (PEI, Polysciences, Cat. #23966) transfection protocol. Env-pseudotyped HIV_DFII_ stocks were produced by co-transfecting plasmids encoding HIV_DFII_ and a plasmid encoding HIV-1 dual-tropic envelope (pSVIII-92HT593.1, NIH HIV Reagent Program, Cat. #3077) at a ratio of 3:1 into HEK293T cells at 50-60% confluency grown in 175 cm^2^ culture flasks. Each flask was transfected with a total amount of 30 μg DNA. The transfection mix was prepared in 2 ml Opti-MEM (ThermoFisher Scientific, Cat. #31985062) as follows: DNA plasmids were diluted in Opti-MEM first, then PEI was added at a ratio of 3:1 PEI:DNA (90 μg PEI). The transfection mix was vortexed for 15 sec and incubated for 15 min at RT. Culture medium was replaced with 20 ml fresh DMEM + 10% FBS <u>without P/S</u>, and 2 ml transfection mix was added to each flask. 16h post transfection, P/S-free medium was replaced with standard culture medium (DMEM+/+), and cells were incubated for another 24h at 37°C, 5% CO_2_. For replication competent HIV NL4-3 Luciferase (a kind gift from Dr. Warner Greene), lentiviral vectors were introduced by Fugene HD transfection (Promega, Cat. #E2311) according to the manufacturer protocols. Cell supernatants were collected 48h post transfection, centrifuged at 4°C for 10 min at 4000 rpm (∼ 3390 x g) and subsequently filtered using 0.22 µm membrane vacuum filter units (MilliporeSigma, Cat. #SCGP00525) to remove cell debris. Virus preparations were concentrated by ultracentrifugation at 20,000 rpm (∼ 50,000 x g) for 2h at 4°C and resuspended in complete media for subsequent storage at −80°C. Virus concentration was estimated by p24 titration (HIV-1 alliance p24 ELISA kit, PerkinElmer, Cat. #NEK050001KT).

### Generation of CRISPRi cell lines

Stable, knockdown cell lines were generated by lentiviral transduction of dCas9-KRAB, as well as single guide RNAs (sgRNAs). We first generated a CRISPRi cell line by transducing J-Lat A72 with the vector lenti-EF1a-dCas9-KRAB-Puro (Addgene #99372, a gift from Kristen Brennand). After selecting with puromycin-containing media (1 μg/ml) for at least one week, J-Lat A72 CRISPRi were tested by qPCR for transgene expression using Cas9-specific primers. Then, J-Lat A72 CRISPRi cells were transduced with the lentiviral vector pLKO5.sgRNA.EFS.tRFP (Addgene #57823, a gift from Benjamin Ebert), containing target-specific sgRNAs (J-Lat A72 CRISPRi CD73KD) or a non-targeting control sgRNA (J-Lat A72 CRISPRi NT). SgRNA design was conducted using the CRISPick-tool developed by the Broad Institute^85^ and sgRNA cloning was performed following the detailed protocol provided by Heckl et al. (2014)^86^. Primer sequences for qPCR and sgRNA cloning are given below.

### J-Lat cell latency reversal

J-Lat 5A8 cells (seeded at 1×10^6^ cells/ml) were incubated with CGS21680 (Sigma-Aldrich, Cat. #C141-5MG) or SCH-58261 (Sigma-Aldrich, Cat. #S4568-5MG) at 37°C for 1h at increasing doses in RPMI+/+, followed by stimulation with 20 nM PMA / 1 μM Ionomycin (PMA/I, Sigma-Aldrich, Cat. #10634-1MG and Cat. #10634-1MG). Untreated cells or cells treated with 0.5% DMSO (Sigma-Aldrich, Cat. #D2650-100ML) served as negative controls. 7h after PMA/I reactivation, cells were washed 2x with PBS (ThermoFisher, Scientific, Cat. #14190-250) and viral transcriptional activity, reflected by GFP expression was measured using LSR II flow cytometer (BD Biosciences).

Analyses of latency reversal in J-Lat A72 CRISPRi CD73KD and J-Lat A72 CRISPRi NT included an additional pre-gating for-RFP positive cells to specifically select for KD cells prior to the quantification of GFP expression.

### Leucocyte isolation and primary CD4+ T cell culture

Peripheral blood mononuclear cells (PBMCs) from HIV-seronegative donors (Vitalant) were isolated by Ficoll-Hypaque density gradient centrifugation (Corning, Inc., Cat. #25-072-Cl) at 2000 rpm (∼ 850 x g) at RT for 30 min, <u>without brake</u>. PBMCs were immediately processed to isolate CD4+ T cells by negative selection using the EasySep Human CD4+ T Cell Isolation Cocktail (StemCell Technologies, Cat. #17952) according to manufacturer’s protocol. Purified CD4+ T cells were cultured in RPMI+/+.

### CD4+ T cell culture under hypoxic conditions

Following CD4+ T cell isolation, cells were activated with with αCD3/αCD28 activating beads (ThermoFisher Scientific, Cat. #11132D) for 48h at a concentration of 1 bead/cell in the presence of 100 U/ml IL-2 (PeproTech, Inc., Cat. #200-02) in RPMI +/+ and were subsequently cultured under hypoxia (HOX - 1% O_2_; Ruskinn Invivo 400 workstation), standard conditions (normoxia - NOX - 21% O_2_), or were treated with 500 μM DMOG (Sigma-Aldrich, Cat. #D3695) up to 5 days post stimulation.

### CD4+ T cell *in vitro* infection

CD4+ T cells were stimulated as described above for 3 days (initial seeding concentration 1×10^6^ cells/ml). At the day of infection, cells were spinoculated in 96-well V-bottom plates (MilliporeSigma, Cat. #M9686-100EA) in 50 μl RPMI+/+ with 100 ng (HIV_DFII_) of p24 per 1×10^6^ cells with 5×10^6^ cells total per well for 2 h at 2350 rpm (1173 × g) at 37°C. After spinoculation, all cells were returned to culture in the presence of 30 U/ml IL-2. Pre-stimulated CD4+ T cells stayed in αCD3/αCD28 activating beads during spininfection and subsequent cell culture. For experiments mimicking hypoxic conditions, cells were treated with 500 μM DMOG (Sigma-Aldrich, Cat. #D3695) or mock-treated with 0.5% DMSO two days after αCD3/αCD28 bead stimulation and 24h before HIV_DFII_ spininfection. Cells were kept in DMOG containing RPMI+/+, in presence of activation beads and IL-2 until sample collection 4 days post infection.

### CD4+ T cell *in vitro* infection and latency reversal

Initially, CD4+ T cells were isolated from peripheral blood as described above and subjected to FACS of CD73+ and CD73-cells. To that aim, cells were stained with APC anti-human CD73 (BioLegend, Cat. #344006) diluted in PBS (1:20) in 100 μl final volume for 20 min at RT. Cells were then washed twice with PBS and resuspended in 500 μl – 1000 μl PBS to achieve high cell concentrations (20 - 40×10^6^ cells/ml) for FACS. Cells were sorted into 15 ml conical tubes containing 1.5 ml RPMI+/+. Cells were cultured for 24h, then infected and rested in the presence of ART to establish *in vitro* latency^47^. Briefly, 100 ng of purified NL4-3-Luciferase was added per 1×10^5^ sorted cells, which were then infected by spinoculation as described above. 24h post virus exposure, 5 µM saquinavir (protease inhibitor, Sigma-Aldrich, Cat. #S8451-50MG) was added to the cell cultures to suppress spreading infection. 5 days later, cells were stimulated with αCD3/αCD28 beads (or left untreated) in the presence of 30 µM raltegravir (integrase inhibitor, Sigma-Aldrich, Cat. #CDS023737-25MG) to prevent new infections. 24h after stimulation, luciferase activity was quantified using the bright glo luciferase assay system (Promega, Cat. #E2610).

### CD4+ T cell staining and processing

Freshly isolated CD4+ T cells were stained for viability, using the fixable Zombie viability dye (1:100; BioLegend, Cat. #423113) according to the manufacturer’s protocol. Subsequently, antibodies APC anti-Human CD73 (1:50), APC-Cy7 anti-Human CD25 (1:100, BD Pharmingen Cat. #557753), and V450 anti-Human CD69 (1:100, BD Horizon Cat. #560740) for cell surface staining diluted in PBS were added and incubated for 20 min at RT. For flow cytometry, cells were washed and fixed in 1% paraformaldehyde (PFA) in PBS after the staining. FACS experiments to isolate populations of interest were performed with live, unfixed cell samples. Flow cytometry analyses were performed on the LSR II flow cytometer (BD Biosciences) or MA900 Multi-Application Cell Sorter (Sony Biotechnologies). All fluorescent-based sorts were conducted on the latter instrument.

### Immunophenotyping of CD4+ T cells from HIV-infected individuals

30 mLs of fresh blood were obtained from each of six HIV-positive individuals on long-term continuous ART (greater than one year) who were stably suppressed (HIV RNA <40 copies/mL); samples were processed as described above. Purified CD4+ T cells were then stained using a panel of antibodies against previously described reservoir markers (see Table 2), as well as CD4, CD73, CD39 and a live/dead staining (ThermoFisher Cat. #L34961). Finally, cells were subjected to flow cytometry measurement on a Cytek Aurora (5L) instrument.

**Table 1:**
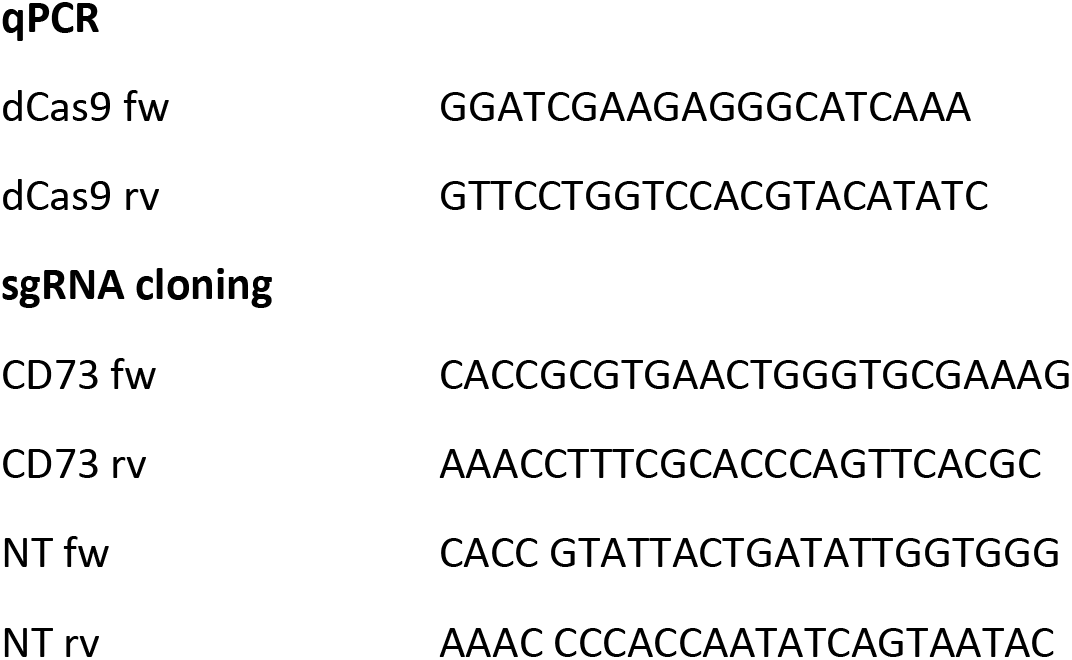
Primers for the generation of CRISPRi cell lines.

**Table 2:** Antibody panel used to phenotype CD4+ T cells from HIV-infected individuals.

| Specificity | fluorophore | Manufacturer | Cat. # |
| --- | --- | --- | --- |
| CD4 | BUV395 | BD Biosciences | 564724 |
| CD30 | BV421 | BD Biosciences | 566253 |
| CD32 | PerCP/Cy5.5 | BioLegend | 303215 |
| CD20 | BV605 | BioLegend | 302333 |
| CD2 | BV510 | BioLegend | 300217 |
| CD279 (PD-1) | PE | BioLegend | 329905 |
| CD49d | BUV737 | BD Biosciences | 612850 |
| CD39 | PE-Cy7 | BioLegend | 329905 |
| CD98 | FITC | BioLegend | 315603 |
| CTLA-4 | BV785 | BioLegend | 369623 |

### Flow cytometry gating and data analysis

Data were analyzed and visualized using the FlowJo software (v.10.7.1). Crosstalk compensations was performed using single-stained samples for each of the fluorochromes and isotype controls were employed to assess antigen positivity and enable specific gating. FACS and flow cytometry gating was performed as follows: low/negative cells for live/dead dyes were selected for single cells and subsequent FSC/SSC scatter plots. Then, antibody gates were defined based on suited /single stained and isotype controls. Gating of HIV_DFII_ reporter expression was based on non-infected, mock-treated (*in vitro* activated) negative control samples to account for activation-dependent increase of cellular background fluorescence.

### Expression Profiling via NanoString

Quantitative RNA and protein expression data were generated using the nCounter Vantage 3D RNA:Protein Immune Cell Profiling Assay (NanoString Technologies, Inc., Cat. #121100019) and the nCounter SPRINT profiler (NanoString Technologies), comprising 770 RNA and 30 protein targets as well as positive and negative controls. 100,000 viable, sorted cells were used per sample, which were processed according to the manufacturer’s instructions. RNA and protein expression values were normalized and analyzed using the nSolver Analysis Software 4.0 and the add-on Advanced Analysis Software 2.0.115 (NanoString Technologies). Samples that did not pass the default control performance and quality parameters were excluded from subsequent analysis. Normalization genes for each sample were automatically selected by the software based on the geNorm algorithm. Biological replicates were grouped according to sample type and the differential expression of each analyte-type (RNA or protein target) was determined in cross-comparisons among all sample types by considering inter-donor differences as confounding variable unless otherwise stated. Intersections of significant targets of individual differential expression analyses were visualized in a Venn diagram using an open source platform from Bioinformatics & Evolutionary Genomics^87^.

Based on the differential expression of each gene, gene sets pre-defined by nanoString, representing different pathways included in this assay, were analyzed by calculating global significance scores for each gene set within each sample as follows: <u>undirected</u> global significance 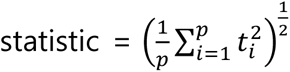, where ti is the t-statistic from the ith pathway gene. The <u>directed</u> global significance statistic is similar to the undirected global significance statistic, but rather than measuring the tendency of a pathway to have differentially expressed genes, it measures the tendency to have over- or under-expressed genes. It is calculated similarly to the undirected global significance score, but it takes the sign of the t-statistics into account: <u>Directed</u> global significance statistic = *sign*(*U*)|(*U*)|^1/2^ where 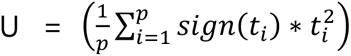 and where sign(U) equals −1 if U is negative and 1 if U is positive (MAN-10030-02^88^).

### CyTOF samples preparation and analysis

For live/dead discrimination, 0.1-1 million cells per sample were treated with cisplatin (Sigma-Aldrich) and fixed with paraformaldehyde (PFA) as previously described^9,28,29^. Briefly, Cells were washed once with contaminant-free PBS (Rockland) with 2 mM EDTA (Corning), centrifuged and resuspend with 25 μM cisplatin in 4 ml PBS/EDTA and incubated for 60 seconds at room temperature, and then quenched with CyFACS (metal contaminant-free PBS supplemented with 0.1% bovine serum albumin and 0.1% sodium azide). Cells were then centrifuged, fixed with 2% PFA in metal contaminant-free PBS and washed 3x with CyFACS. These fixed cells were stored at −80°C until CyTOF staining.

Prior to CyTOF staining, multiple samples were barcoded using the Cell-ID 20-Plex Pd Barcoding Kit according to manufacturer’s instructions (Fluidigm). Briefly, each sample was washed twice with Barcode Perm buffer (Fluidigm), and incubated for 30 min with the appropriate barcode at a 1:90 ratio. Cells were then washed with 0.8 ml Maxpar Cell Staining buffer (Fluidigm) in Nunc^TM^ 96 Deep-Well polystyrene plates (Thermo Fisher), followed by CyFACS. Barcoded samples were combined and blocked with sera from mouse (Thermo Fisher), rat (Thermo Fisher), and human (AB serum, Sigma-Aldrich) for 15 minutes at 4°C. Cells were washed twice with CyFACS buffer and stained with the cocktail of CyTOF surface staining antibodies (Table S1) for 45 min on ice. Subsequently, cells were washed 3X with CyFACS buffer and fixed overnight at 4°C with 2% PFA (Electron Microscopy Sciences) in metal contaminant-free PBS. For intracellular staining, cells were permeabilized by incubation with fix/perm buffer (eBioscience) for 30 min at 4°C and washed twice with Permeabilization Buffer (eBioscience). Cells were blocked again with sera from mouse and rat for 15 min on ice, washed twice with Permeabilization Buffer (eBioscience), and stained with the cocktail of CyTOF intracellular staining antibodies (Table S1) for 45 min on ice. Cells were then washed once with CyFACS and incubated for 20 min at room temperature with 250 nM Cell-IDTM DNA Intercalator-Ir (Fluidigm) in 2% PFA diluted in PBS. Cells were washed twice with CyFACS, once with Maxpar Cell Staining Buffer (Fluidigm), once with Maxpar PBS (Fluidigm), and once with Maxpar Cell Acquisition Solution (CAS, Fluidigm). Immediately prior to acquisition, cells were resuspended in EQTM calibration beads (Fluidigm) diluted in CAS. Cells were acquired at a rate of 250-350 events/sec on a CyTOF2 instrument (Fluidigm) at the UCSF Parnassus single cell analysis facility.

Data were normalized to EQTM calibration beads and de-barcoded with CyTOF software (Fluidigm). Normalized data were imported into FlowJo (BD) for gating (cell, intact, live, single events) and heatmap was generated in Cytobank.

### RNA Sequencing

Freshly isolated CD4+ T cells from healthy blood donors were sorted based on their CD73 expression as described above. 3×10^6^ cells were collected per sample and stored as dry cell pellets at −80 °C. RNA preparation, library preparation and mRNA sequencing were conducted at Genewiz (USA). Paired-end sequencing was performed using the Illumina NovaSeq 6000 instrument to obtain a minimum of 20 million read pairs per sample with a read length of 2×150 bp. Sequence reads were trimmed to remove possible adapter sequences and nucleotides with poor quality using Trimmomatic v.0.36. The trimmed reads were mapped to the Homo sapiens GRCh38 reference genome available on ENSEMBL using the STAR aligner v.2.5.2b. Unique gene hit counts were calculated by using featureCounts from the Subread package v.1.5.2. The hit counts were summarized and reported using the gene_id feature in the annotation file. Only unique reads that fell within exon regions were counted. After extraction of gene hit counts, the gene hit counts table was used for downstream differential expression analysis. Using DESeq2, a comparison of gene expression between samples was performed adjusting for the donor effect as confounding variable. The Wald test was used to generate p-values and log_2_ fold changes. Genes with an adjusted p-value < 0.05 (Benjamini-Hochberg method) and absolute log_2_ fold change > 1 were called as differentially expressed genes for each comparison. A gene ontology analysis was performed on the statistically significant set of genes by implementing the software GeneSCF v.1.1-p2. The goa_human GO list was used to cluster the set of genes based on their biological processes and determine their statistical significance. A list of genes clustered based on their gene ontologies was generated.

### *In Situ* Detection of HIV and Cellular Markers

The experimental procedure for the in situ and immunofluorescence staining has been described in detail in previous publications^52,53^. The assay enables the detection of HIV-integrated DNA, viral mRNA, viral proteins, and several cellular markers in the same assay. Sample preparation, data acquisition and subsequent analyses were conducted in the laboratory of Dr. Eliseo Eugenin (UTMB, Texas).

### Tissue samples

Tissues from ART-suppressed individuals who have been on treatment for at least 6 months and had viral loads below clinical detection limits (<50 RNA copies/ml), as well as tissues from HIV-negative and ART-naïve viremic individuals with high plasma loads (>50 RNA copies/ml) were part of an ongoing research protocol approved by University of Texas Medical Branch (UTMB). Further clinical data and additional information are available and will be provided upon request by the lead contact Eliseo Eugenin. All tissues were obtained with full, written consent from the study participants and freshly collected specimens were immediately fixed with 4% PFA, then mounted into paraffin blocks, subjected to tissue sectioning, and ultimately to ab analysis by *in situ* and immunostaining.

### Staining procedures

Paraffin-embedded slides containing the tissue samples were consecutively immersed in the following solutions: xylene for 5 min (2 times), 100% EtOH for 3 min, 100% EtOH for 3 min, 95% EtOH for 3 min, 90% EtOH for 3 min, 70% EtOH for 3 min, 60% EtOH for 3 min, 50% EtOH for 3 min, miliQ H_2_O for 3 min. Then, tissue was encircled with ImmEdge Pen to reduce the reagent volume needed to cover the specimens. Finally, slides were immersed in miliQ H_2_O for 3 min. <u>For Protein K treatment</u>, tissues were incubated with proteinase K diluted 1:10 in 1X TBS (Fisher Scientific, Cat. #BP24711; and PNA ISH kit, Agilent Dako, Cat. #K5201) for 10 min at RT in a humidity chamber. Next, slides were immersed in miliQ H2O for 3 min, then immersed in 95% EtOH for 20 sec and finally, the slides were left air-dry for 5 min. <u>For HIV DNA probe hybridization</u> tissues were incubated with 10 µM PNA DNA probe for Nef-PNA Alexa Fluor 488 and Alu-PNA Cy5 (PNA Bio). Next, slides were placed in a pre-warmed humidity chamber and incubated at 42°C for 30 min, then the temperature was raised to 55°C for an additional 1 h incubation. Subsequently, tissues were incubated using Preheat Stringent Wash working solution (PNA ISH kit) diluted 1:60 in 1X TBS for 25 min in an orbital shaker at 55°C. Slides were equilibrated to RT by brief immersion in TBS for 20 sec. <u>HIV</u> <u>mRNA detection</u> followed the manufacturer’s protocol for RNAscope 2.5 HD Detection Reagent-RED (Advanced Cell Diagnostics, Inc., Cat. #322360). Probe for HIV Gag-pol was added to the tissue samples and incubated for 30 min at 42°C and then 50 min at 55°C. Next, samples were incubated in Preheat Stringent Wash working solution diluted 1:60 in 1X TBS (PNA ISH kit) for 15 min in an orbital shaker at 55°C. Finally, slides were immersed in 1X TBS for 20 sec. <u>For HIV or cellular</u> <u>protein detection,</u> antigen retrieval was performed by incubating slide sections in commercial antigen retrieval solution (Agilent Dako, Cat. #S1700) for 30 min in a water-bath at 80°C. Next, slides were removed from the bath and cooled down in 1X TBS. Samples were permeabilized with 0.1% Triton X-100 (Sigma-Aldrich, Cat. #X100) for 2 min and then washed in 1X TBS for 5 min three times. Unspecific antibody binding sites were blocked by incubating samples with a freshly prepared blocking solution. Afterwards, sections were incubated overnight at 4°C using a humidity chamber (10 ml of blocking solution: 1 mL 0.5 M EDTA, 100 ul Fish Gelatin from cold water 45%, 0.1 g Albumin from Bovine serum Fraction V, 100 ul horse serum, 5% human serum, 9 mL miliQ H_2_O). Anti-p24 primary antibodies were added to the samples diluted in a blocking solution and incubated at 4°C overnight. To perform these analyses, large amounts of antibodies (obtained from the NIH AIDS repository) were purified and pre-absorbed in uninfected human tissues as well as tested in cell lines to assure specificity and proper binding to the target (see Donoso et al.^52^ for details). Following antibody staining, slides were washed in 1X TBS 5 min for three times to eliminate unbound antibodies. Secondary antibodies were added at the appropriate dilutions and incubated for 2h at RT. Slides were washed three times in 1X TBS for 5 min to eliminate unbound antibodies. Next, slides were mounted using Prolong Diamond Antifade Mount medium containing DAPI (ThermoFisher Scientific, Cat. #P36931). Slides were kept in the dark at 4°C.

### Image acquisition and analysis

Cells were examined by confocal microscopy using an A1 Nikon confocal microscope with spectral detection and unmixing systems. Image analysis was performed using the Nikon NIS Elements Advanced Research imaging software (Nikon Instruments, Japan). The automated image segmentation and analysis are based on the following premises: For detection of HIV-integrated DNA, <u>first</u>, automatic or manual detection of cells that are positive for HIV-DNA and <u>second</u>, the HIV-DNA probe has to colocalize with DAPI and Alu repeats staining with a Pearson’s correlation coefficient of at least 0.8 as described previously^89^. These two conditions are essential for HIV-integrated DNA, or the signal is considered negative or unspecific. For detection of HIV-mRNA, <u>first</u>, low colocalization with DAPI or Alu-repeats (0.2 Pearson’s correlation coefficient or below) and <u>second</u>, presence in cells with HIV-DNA signal. The sensitivity, accuracy and specificity of the system were previously validated in the laboratory of our collaborator in two well characterized T cell lines A3.01 (uninfected) and ACH-2 (HIV-infected) and two monocytic cell lines, HL-60 (uninfected) and OM-10 (HIV-infected).

To analyze local microenvironments of CD3+ cells, CD73 expression was assessed in cell neighborhoods by lineplot analysis. Image segmentation and quantification were performed using Nikon NIS Elements Advanced Research imaging software (Nikon Instruments, Japan) and Ilastik^90^. CD73 signals were analyzed along straight lines of 80 µM length, placing the zero point at the center of the line, at the position of the respective CD3-positive cell. Obtained CD73 intensities were normalized to the mean CD73 intensity in CD3+ HIV-DNA-/p24+ (ART suppr.) samples (see Figure S7).

### Quantification and Statistical Analysis

Statistical details are given in the figure legends. All statistical analyses were performed using GraphPad Prism software version 9. P-values ≤ 0.05 were considered statistically significant. A Student’s two-tailed t-test was used for two-way column analyses. ANOVA tests were used for multiple comparisons. P-values are denoted in the figure panels. Data are presented as means with error bars indicating the standard error of the mean (SEM) unless otherwise stated.

## DATA AVAILABILITY

The datasets generated and/or analyzed during the current study are available from the corresponding author on reasonable request.

## ACKNOWLEDGEMENTS

The authors would like to acknowledge Guorui Xie, PhD and Ashley George, PhD for helpful input regarding the CyTOF analysis. The authors thank Leonard Chavez, PhD and Shivani Desai for providing reagents, Vivienne Schneider for experimental support, as well as Konstantinos Georgiou, PhD and Zain Y. Dossani, PhD for guidance and support in data analysis. The authors sincerely thank Rebecca Hoh for her help in obtaining clinical specimens for our study.

This study was supported by National Institutes of Health (NIH) grants R01AI150449 and R01MH112457 (to S.K.P.). We would like to thank The National NeuroAIDS Tissue Consortium (NNTC) for providing all human samples and associated information. The NNTC is made possible through funding from the NIMH and NINDS by the following grants: Manhattan HIV Brain Bank (MHBB): U24MH100931; Texas NeuroAIDS Research Center (TNRC): U24MH100930; National Neurological AIDS Bank (NNAB): U24MH100929; California NeuroAIDS Tissue Network (CNTN): U24MH100928; and Data Coordinating Center (DCC); R01AI147777; R01AI127219; UM1AI64567; UM1AI64559. The staining and analysis were funded by the National Institute of Mental Health grant, MH096625 and MH128032 and the National Institute of Neurological Disorders and Stroke, NS105584 (to E. A. E.).

