## Supplement Information for "The Hypoxia-regulated Ectonucleotidase CD73 is a Host Determinant of HIV Latency"

**Table S1: List of CyTOF staining antibodies.**

| Antibody | Metal label | Clone | Vendor |
|---|---|---|---|
| CD49d (□4) | 141Pr | 9F10 | Fluidigm |
| CD19 | 142Nd | HIB19 | Fluidigm |
| CCR5 | 144Nd | NP6G4 | Fluidigm |
| CD8 | 146Nd | RPAT8 | Fluidigm |
| CD7 | 147Sm | CD76B7 | Fluidigm |
| ICOS | 148Nd | C398.4A | Fluidigm |
| CD103 | 151Eu | Ber-ACT8 | Fluidigm |
| CD62L | 153Eu | DREG56 | Fluidigm |
| TIGIT | 154Sm | MBSA43 | Fluidigm |
| CCR6 | 155Gd | G034E3 | In-house |
| CD29 (□1) | 156Gd | TS2/16 | Fluidigm |
| OX40 | 158Gd | ACT35 | Fluidigm |
| CCR7 | 159Tb | G043H7 | Fluidigm |
| CD28 | 160Gd | CD28.2 | Fluidigm |
| CD45RO | 161Dy | UCHL1 | In-house |
| CD69 | 162Dy | FN50 | Fluidigm |
| CRTH2 | 163Dy | BM16 | Fluidigm |
| PD1 | 164Dy | EH12.1 | In-house |
| CD127 | 165Ho | A019D5 | Fluidigm |
| CXCR5 | 166Er | RF8B2 | In-house |
| CD27 | 167Er | L128 | Fluidigm |
| CD30 | 168Er | BerH8 | In-house |
| CD45RA | 169Tm | HI100 | Fluidigm |
| CD3 | 170Er | UCHT1 | Fluidigm |
| CD57 | 171Yb | HCD57 | In-house |
| CD38 | 172Yb | HIT2 | Fluidigm |
| α4β7 | 173Yb | Act1 | In-house |
| CD4 | 174Yb | SK3 | Fluidigm |
| CXCR4 | 175Lu | 12G5 | Fluidigm |
| CD25 | 176Yb | M-A251 | In-house |
| HLADR | 112Cd | Tu36 | Invitrogen |
| ROR□t[#] | 115Di | AFKJS-9 | In-house |

| | | | |
|---|---|---|---|
| NFAT1# | 143Nd | D43B1 | Fluidigm |
| BIRC5# | 145Nd | 91630 | In-house |
| Tbet# | 149Sm | eBio4B10 (4B10) | In-house |
| KC57# | 150Nd | KC57 | In-house |
| Blimp1# | 152Sm | 6D3 | In-house |
| CTLA4# | 157Gd | 14D3 | In-house |
| KC57# | 209Bi | KC57 | In-house |

#*Intracellular antibodies*

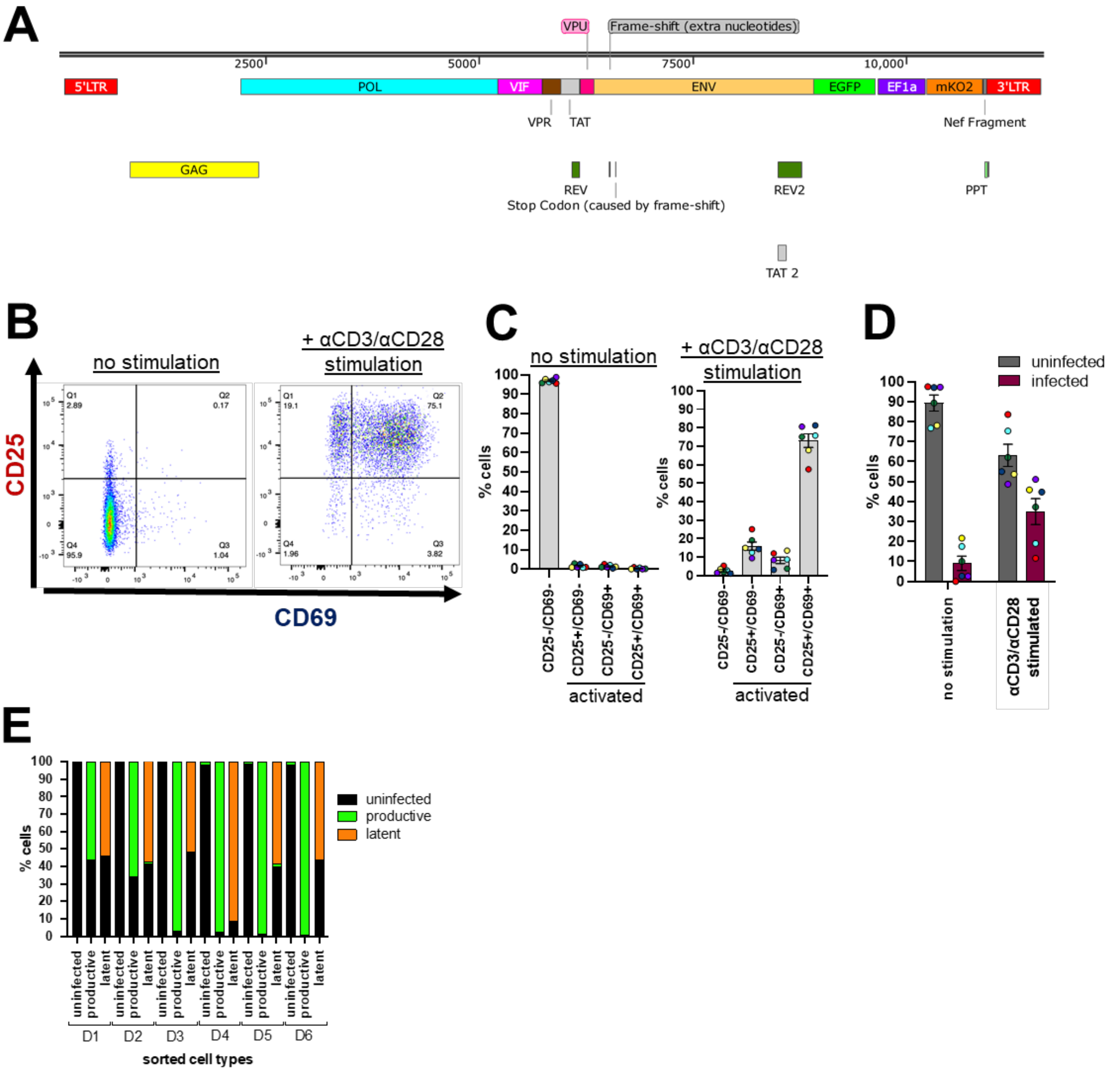


**Figure S1: Development of an experimental workflow for expression profiling of HIV-infected cells.** (A) Genomic organization of the dual-color reporter virus $HIV_{DFII}$. $HIV_{DFII}$ was adapted from the previous dual-color HIV reporter version HIV Duo-Fluo I ($HIV_{DFI}$)[26,,27]. The latent reporter mCherry in $HIV_{DFI}$ was exchanged by molecular cloning with the brighter fluorescent protein mKO2 to improve detection of latently-infected cells. The virus harbors a non-sense mutation in the ORF of *env*, limiting infections to a single round. This recombinant virus encodes for two separate fluorescent markers with enhanced green fluorescent protein (eGFP) indicating active viral transcription and mKusabira-Orange2 (mKO2) reporting viral integration. Cells expressing eGFP alone or eGFP and mKO2 are considered productively-infected; cells expressing mKO2 alone are considered latently-infected; cells lacking the expression of either reporter are considered uninfected. (B) The frequency of live, single, activated CD4+ T cells was determined based on activation markers CD69 (early activation) and CD25 (late activation). Cells were either left untreated or stimulated with αCD3/αCD28 beads for three days and subsequently infected with $HIV_{DFII}$. $1x10^6$ cells were then stained with anti-CD25 and anti-CD69 fluorescent antibodies to measure surface expression using flow cytometry. (B) Shown are representative flow plots from one donor for CD25/CD69 surface expression in unstimulated and αCD3/αCD28 stimulated cells before $HIV_{DFII}$ infection. (C) The frequency of CD25/CD69 expressing cells was quantified three days post stimulation for 6 donors. (D) Data represent the frequency of $HIV_{DFII}$-infected cells left untreated or stimulated with αCD3/αCD28 beads measured for 6 donors four days after $HIV_{DFII}$ infection. (E) The purity of all sorted samples was assessed post processing by remeasuring sample aliquots using flow cytometry. Data show the percentage of cells composing the sorted samples for each donor normalized to the total gated events.

**Table S2: Targets identified in the differential expression analyses comparing latently-infected cells to both other sorted samples with a p-value cutoff at $p < 0.05$.** Targets are sorted by p-value of latent vs productive comparison.

| | latent vs uninfected | | latent vs productive | |
|---|---|---|---|---|
| Target | log2 fold change | p-value | log2 fold change | p-value |
| **IL23A-mRNA** | 0.483 | 0.00365 | 0.81 | 0.0000954 |
| **CD47-mRNA** | -0.106 | 0.0151 | -0.221 | 0.000114 |
| **NT5E-protein** | 0.644 | 0.000111 | 0.638 | 0.000119 |
| **IL8-mRNA** | 0.785 | 0.0114 | 1.4 | 0.000295 |
| **CASP1-mRNA** | -0.166 | 0.0434 | -0.361 | 0.000519 |
| **FOS-mRNA** | 2.56 | 3.18E-05 | 1.53 | 1.57E-03 |
| **CCR4-mRNA** | -0.15 | 0.032 | -0.237 | 0.0028 |
| **NFKBIA-mRNA** | 0.39 | 0.00118 | 0.342 | 0.00281 |
| **STAT1-mRNA** | -0.12 | 0.0326 | -0.19 | 0.00286 |
| **CD53-mRNA** | -0.145 | 0.0132 | -0.184 | 0.00345 |
| **ITGAX-mRNA** | 0.282 | 0.00267 | 0.263 | 0.00408 |
| **IL24-mRNA** | 0.705 | 0.00152 | 0.592 | 0.00466 |
| **TNFAIP3-mRNA** | 0.321 | 0.000206 | 0.203 | 0.00492 |
| **TAPBP-mRNA** | -0.16 | 0.00714 | -0.168 | 0.00534 |
| **GNLY-mRNA** | -0.303 | 0.0441 | 0.431 | 0.0104 |
| **HLA-DRA-mRNA** | -0.46 | 0.00119 | -0.318 | 0.0114 |
| **CCL5-mRNA** | 0.286 | 0.000383 | 0.169 | 0.0114 |
| **CD2-mRNA** | 0.238 | 0.0174 | 0.252 | 0.0133 |
| **TP53-mRNA** | -0.151 | 0.0134 | -0.143 | 0.0174 |
| **HLA-DRB3-mRNA** | -0.325 | 0.0273 | -0.356 | 0.018 |
| **ENTPD1-mRNA** | -0.529 | 0.0109 | -0.467 | 0.0209 |
| **CSF2-mRNA** | 0.577 | 0.0157 | 0.528 | 0.0241 |
| **MIF-mRNA** | 0.101 | 0.0189 | 0.0942 | 0.0258 |
| **CD83-mRNA** | 0.33 | 0.0188 | 0.284 | 0.0361 |
| **CYLD-mRNA** | -0.165 | 0.032 | -0.156 | 0.0404 |
| **TIGIT-mRNA** | -0.335 | 0.0387 | -0.328 | 0.0424 |
| **IL1RAP-mRNA** | 0.308 | 0.00238 | 0.175 | 0.0451 |

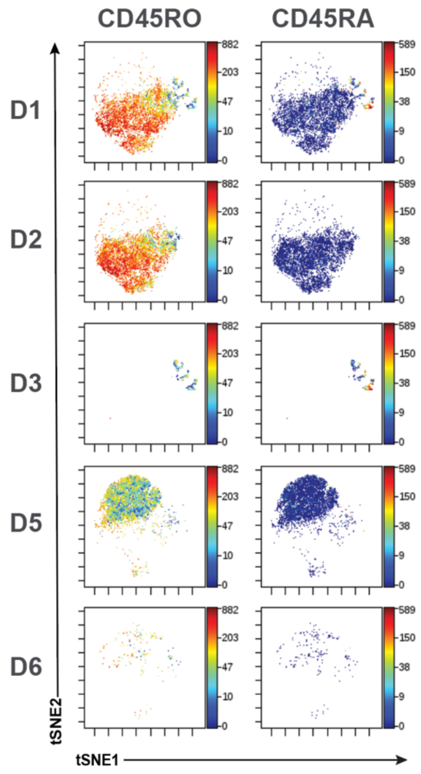


**Figure S2: Donor-dependent differences mainly drive cell segregation patterns.** Displayed are individual tSNE plots for each donor. tSNEs are color-coded according to expression levels of the antigen listed at the top (with red corresponding to highest expression, and blue the lowest). Strong donor-to-donor differences are evident by the apparent segregation of the various donors in the tSNE space.

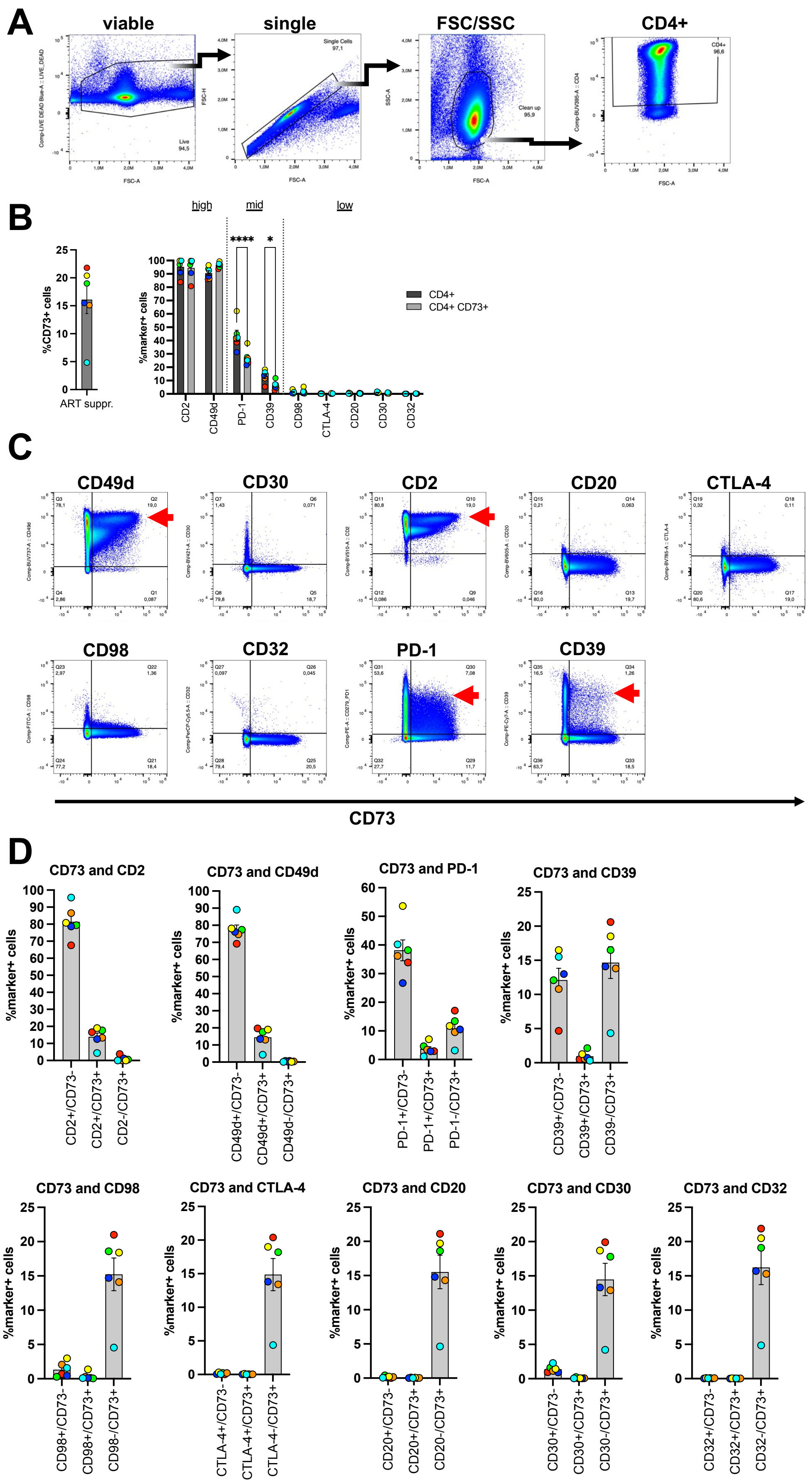

A
viable
single
FSC/SSC
CD4+
Live 94,5
Single Cells 97,1
Clean up 95,9
CD4+ 96,6
FSC-A
FSC-H
SSC-A
B
%CD73+ cells
ART suppr.
high
mid
low
****
*
%marker+ cells
CD4+
CD4+ CD73+
CD2
CD49d
PD-1
CD39
CD98
CTLA-4
CD20
CD30
CD32
C
CD49d
CD30
CD2
CD20
CTLA-4
CD98
CD32
PD-1
CD39
CD73
D
CD73 and CD2
CD73 and CD49d
CD73 and PD-1
CD73 and CD39
CD73 and CD98
CD73 and CTLA-4
CD73 and CD20
CD73 and CD30
CD73 and CD32
%marker+ cells
CD2+/CD73-
CD2+/CD73+
CD2-/CD73+
CD49d+/CD73-
CD49d+/CD73+
CD49d-/CD73+
PD-1+/CD73-
PD-1+/CD73+
PD-1-/CD73+
CD39+/CD73-
CD39+/CD73+
CD39-/CD73+
CD98+/CD73-
CD98+/CD73+
CD98-/CD73+
CTLA-4+/CD73-
CTLA-4+/CD73+
CTLA-4-/CD73+
CD20+/CD73-
CD20+/CD73+
CD20-/CD73+
CD30+/CD73-
CD30+/CD73+
CD30-/CD73+
CD32+/CD73-
CD32+/CD73+
CD32-/CD73+

**Figure S3: CD73 is highly expressed on a subset of PD-1, CD39, Cd49d and CD2 positive cells.** Immunofluorescence staining of CD4+ T cells from six ART-suppressed, HIV-infected individuals was performed, followed by flow cytometry. A flow cytometry panel was designed to investigate the co-expression of CD73 and previously described HIV reservoir markers. (A) Cells were pre-gated for viable (live/dead negative) singlets (single), excluding debris (FSC/SSC) and CD4-negative cells. (B) Frequency of CD73+ cells among CD4+ T cells (left graph) and indicated marker-positive subpopulations among CD4+ and CD4+/CD73+ cells, respectively (right graph). Significance was assessed by ordinary two-way ANOVA, followed by Sidak's multiple comparisons test: **** adj. $p < 0.0001$; * adj. $p = 0.01–0.05$. (C) Flow cytometry plots showing expression levels of the cellular proteins as described in B plotted against CD73. Double positive sub-populations were identified for CD39, PD-1, CD49d and CD2 (high co-expression indicated by red arrows). (D) Quantification of marker expression as shown in C for all 6 participants.

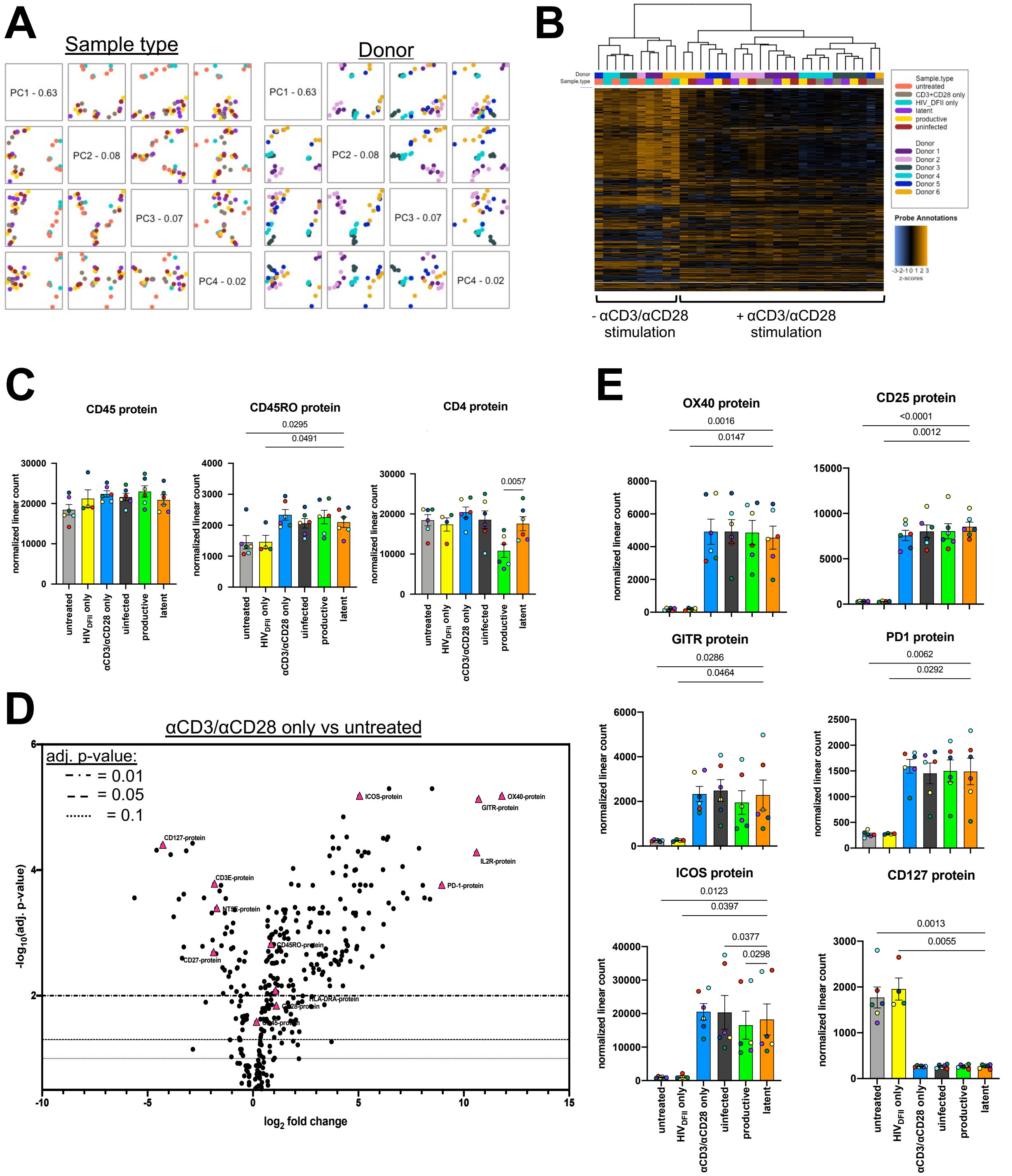


**Figure S4: Response to TCR stimulation dominates expression patterns across samples and donors but does not associate with HIV latency.** (A) Displayed are principal component analyses, which transform and display high-dimensional gene expression data on a 2D plane based on a distance matrix. Each dot represents one sample and is colored by the attributes 'sample type' or 'donor' (color legend on the right). Principal component 1 (PC1) captures the highest level of variance, followed by PC2, PC3 and PC4. Each PC is plotted twice vs the other (image is diagonally mirrored). The diagonal boxes denote the PC name and its contribution to the total variance in percent. Plots in the same row show this specific PC on their y-axis and plots in the same column display this PC on their x-axis. Samples with similar features are clustered together while differing samples are further apart. (B) Heatmap visualizing normalized gene expression data. Each row shows counts of a single probe (target) and each column represents an individual sample. Colored horizontal bars along the top of the heatmap identify assigned sample attributes (see color legend). Hierarchical clustering was used to generate dendrograms. (C) Bar graphs represent the average normalized linear expression levels of generic T cell markers CD45, CD45RO and CD4 in all samples for all 6 donors. (D) Panel specific CD4+ T cell response to TCR stimulation. The volcano plot visualizes the differential expression analysis, comparing normalized target expression in αCD3/αCD28 only samples vs untreated samples of all six donors. Each data point in the scatter plot represents one target species. Protein targets are highlighted as pink triangles. The $\log_2$ fold change of each target is displayed on the x-axis and the -$\log_{10}$ of its adjusted (adj.) p-value (using the Benjamini-Hochberg method) on the y-axis. Horizontal lines indicate adj. p-value thresholds. (E) Bar graphs show the average normalized linear expression levels of defined T cell activation markers for all 6 donors. Colors dots indicate individual donors. Error bars show SEM.

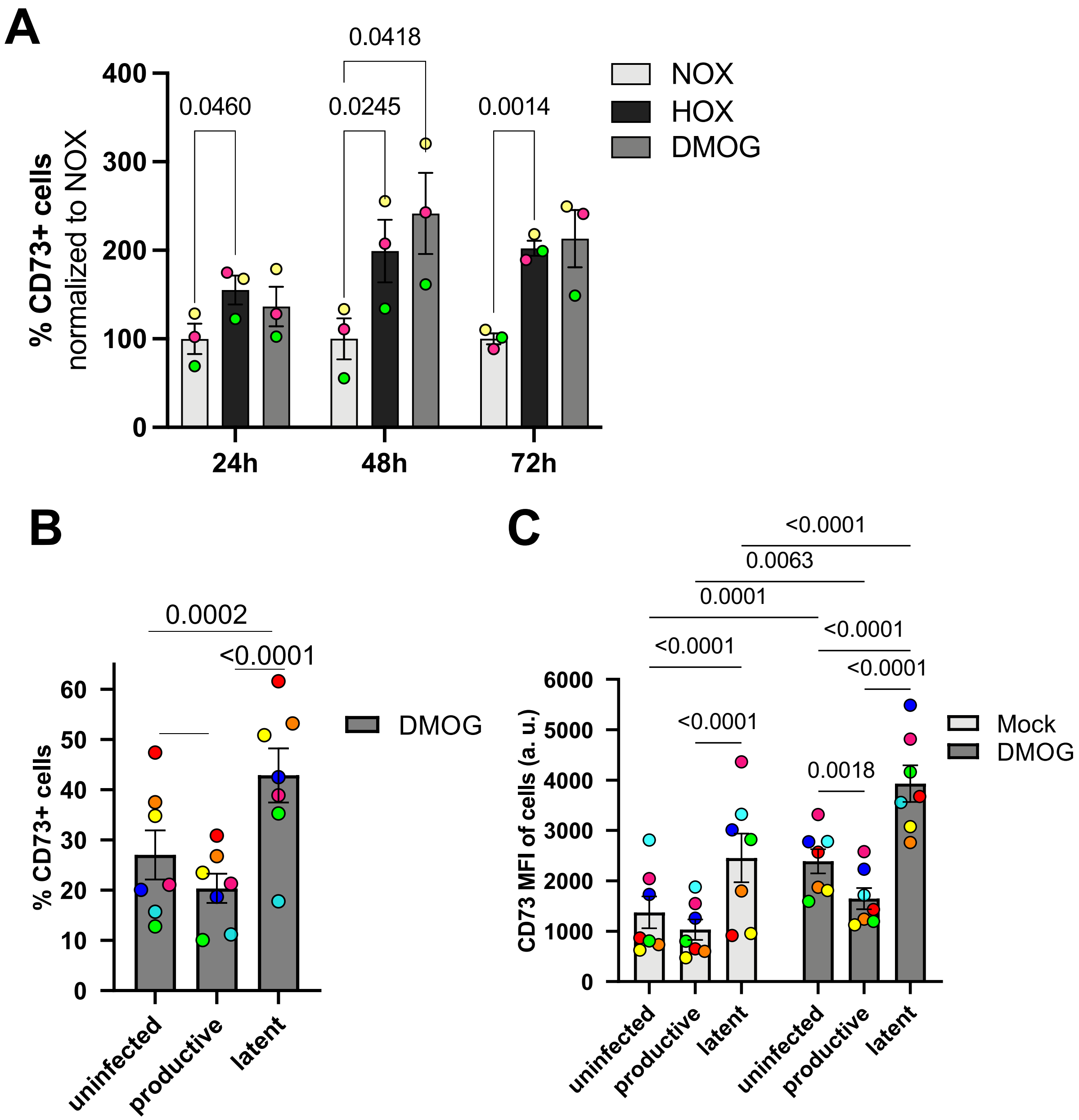


**Figure S5: The enrichment of CD73 surface expression in HIV latently-infected cells is reinforced under hypoxic conditions.** Primary CD4+ T cells were activated with CD3/CD28 activation beads for 48h and subsequently incubated under HOX (1% O2), NOX (21% O2) or NOX with 500 µM DMOG for indicated time points. After the respective time surface CD73 expression was assessed via immunofluorescence staining and flow cytometry. Each bar represents the mean with SEM from 3 biological replicates. Statistical significance was evaulated using two-way ANOVA. At each time point, the percentage of CD73-positive cells was normalized to the respective NOX sample. (B) The Frequency of CD73+ cells and (C) CD73 surface expression per cell reflected by the mean fluorescence intensity (CD73-MFI) is displayed for 7 donors. Colored dots indicate individual donors. P-values displayed within the diagrams were generated by performing a (B) one-way ANOVA with Tukey's multiple comparisons test and (C) a two-way ANOVA with Fisher's LSD. P-value threshold for significance was at $p < 0.05$. Error bars show SEM.

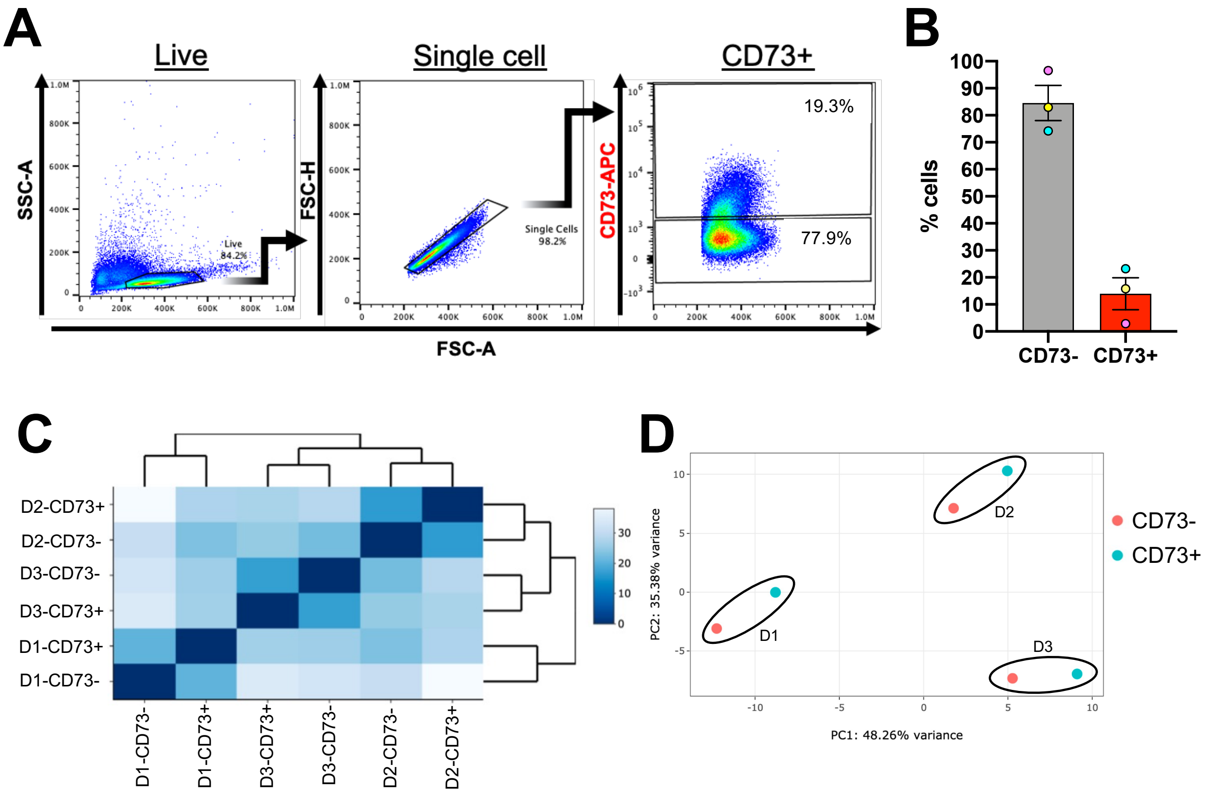


**Figure S6: CD73+ cells exhibit specific immunoregulation and signaling cascade patterns.** (A) Representative flow plots demonstrate the gating strategy for FACS of CD73- and CD73+ CD4+ T cells directly following blood processing, CD4+ T cell isolation and CD73 staining. (B) The frequency of CD4+ CD73+/- T cells was measured for three donors. Error bars show SEM. (C) Heatmap depicts overall similarity among sorted samples assessed by their Euclidean distance. Clustering and dendrogram organization reflect relatedness between samples. (D) PCA plot displaying PC1 and PC2. The percentage of the total variance per direction is shown on the respective axes. D = Donor.

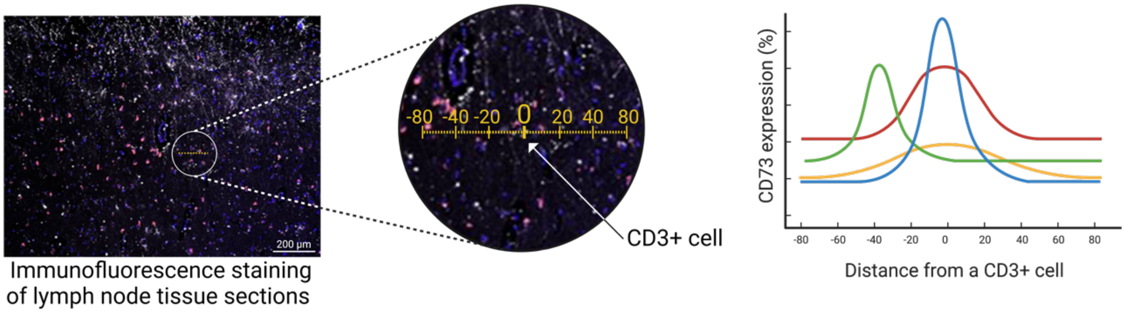


$$\text{CD73 expression (\%)} = 100 \times \frac{\text{CD73 signal}}{\text{mean CD73 (CD3+, HIV DNA}^{-}\text{, p24}^{+}\text{, ART suppressed)}}$$

**Figure S7:** Schematic representing the analysis of CD3+ cell microenvironments. CD3-positive cells were identified in 3D reconstructed images using Ilastik and NIS Elements Advanced Research imaging software. Then, 80 μM lineplots were placed across CD3+ cells, defining the center of the line as the origin of the one-dimensional coordinate system and the position of the CD3+ cell. CD73 intensities were assessed in 10 μM steps and normalized to the mean CD73 intensity in the CD3+, HIV DNA-, p24+, ART suppressed sample. 40-98 images were analyzed per sample.
